# A combined lipidomic and proteomic profiling of *Arabidopsis thaliana* plasma membrane

**DOI:** 10.1101/2023.05.14.540643

**Authors:** Delphine Bahammou, Ghislaine Recorbet, Adiilah Mamode Cassim, Franck Robert, Thierry Balliau, Pierre Van Delft, Youcef Haddad, Sébastien Mongrand, Laetitia Fouillen, Françoise Simon-Plas

**Affiliations:** Laboratoire de Biogenèse Membranaire, CNRS, Université. Bordeaux, (UMR 5200), F-33140 Villenave d’Ornon, France; UMR Agroécologie, INRAE, Institut Agro Dijon, Université Bourgogne Franche-Comté, F-21000 Dijon, France; Université Paris-Saclay, INRAE, CNRS, AgroParisTech, GQE-Le Moulon, PAPPSO, F-91190, Gif-Sur-Yvette, France

## Abstract

The plant plasma membrane (PM) plays a key role in nutrition, cell homeostasis, perception of environmental signals, and set-up of appropriate adaptive responses. An exhaustive and quantitative description of the whole set of lipids and proteins constituting the PM is thus necessary to understand how the way these components, are organized and interact together, allow to fulfill such essential physiological functions. Here we provide by state-of-the-art approaches the first combined reference of the plant PM lipidome and proteome from *Arabidopsis thaliana* suspension cell culture. We identified a reproducible core set of 2,165 proteins (406 of which had not been shown associated to PM previously), which is by far the largest set of available data concerning the plant PM proteome. Using the same samples, we combined lipidomic approaches, allowing the identification and quantification of an unprecedented repertoire of 405 molecular species of lipids. We showed that the different classes of lipids (sterols, phospholipids, and sphingolipids) are present in similar proportions in the plant PM. Within each lipid class, the precise amount of each lipid family and the relative proportion of each molecular species were further determined, allowing us to establish the complete lipidome of Arabidopsis PM, and highlighting specific characteristics of the different molecular species of lipids (for instance fatty acyl chain length and saturation according to the polar head). Results obtained are consistent with the plant PM being an ordered mosaic of domains and point to a finely tuned adjustment of the molecular characteristics of lipids and proteins. More than a hundred proteins related to lipid metabolism, transport or signaling have been identified and put in perspective of the lipids with which they are associated. All these results provide an overall view of both the organization and the functioning of the PM.

## Introduction

In all organisms, the plasma membrane (PM) forms a selective barrier between the cell and the extracellular environment and plays multiple roles, including regulation of nutrient fluxes, perception of cellular environment and intricate orchestration of signal transduction allowing translation of external signals in finely tuned appropriate adaptive responses. Throughout the different kingdoms, the PM of all living cells is made of a bilayer of lipids (with polar heads facing intra and extracellular medium) within which proteins are inserted (1). At the time this model of “fluid mosaic” was established, it was considered that lipids were mainly responsible of forming the hydrophobic barrier preventing the passive transfer of solutes and that proteins were responsible of all the active transport and signaling processes. If the fundamentals of this model have not been questioned, it has been tremendously enriched by the identification of additional elements concerning the organization and functioning of the PM (2), notably through reciprocal interactions between lipids and proteins.

A first element is the role of lipids as direct regulators of protein activity. Indeed, the lipid molecules in contact with an intrinsic membrane protein act as a solvent for the protein: these lipids have been referred to as boundary lipids or as annular lipids, to denote the fact that they form an annular shell of lipid around the protein (3,4). Annular lipids have a slower dynamics than the bulk ones, and provide through their molecular characteristics (charge, steric hindrance of the polar head, length and unsaturation of the acyl chain) a well-suited nano-environment for the surrounded intrinsic protein. Some membrane lipids might also act as cofactors directly regulating protein activity, these lipids often co-purifying with the protein and being resolved in high-resolution structures of the protein (5). The involvement of PIP_2_ in the regulation of integral PM proteins has for instance emerged as a key process (6,7).

Membrane proteins are also quite sensitive to biophysical characteristics of the membrane which are intimately linked to its lipid composition. A typical feature of the membrane is the thickness of the hydrophobic core of the bilayer, which is expected to match the hydrophobic thickness of any protein embedded in the bilayer, since the cost of exposing either fatty acyl chains or hydrophobic amino acids to water is very high (8). Any mismatch between the hydrophobic thickness of the lipid bilayer and the protein would lead to distortion of the lipid bilayer, or the protein, or both, to minimize it, as evidenced by the modulation of membrane protein structure and activity consecutive to changes in fatty acyl chain length (9,10). This points to a necessary fine adjustment between the length of protein TM helices and the thickness of the hydrophobic core of the lipid bilayer determined by the length of lipid acyl chains.

One of the major breakthroughs in the understanding of membrane organization since the description of the fluid mosaic model was the discovery that PM components are not homogeneously distributed within the membrane. The « lipid raft » hypothesis, bringing together structural and functional animal cell membrane organization thus proposed rafts as PM platforms of high molecular order (reflecting lipid packing), enriched in cholesterol and sphingolipids, in which proteins involved in signaling can selectively interact with effector molecules (11). Since an ever-increasing number of proteins and lipids have been described as organized into nanometric membrane compartments, the PM of living cells must now be acknowledged as a dynamic, highly compartmentalized mosaic wherein membrane domains with different biophysical properties co-exist at different scales (12,13). The confining of membrane proteins in distinct nanodomains has been hypothesized as a way for plant cells to use similar proteins to respond differentially to various stimuli (14,15). All studies performed either on artificial membranes, isolated biological membranes or PM of living cells demonstrated unambiguously that lipids were critical regulators of plant PM organization at a nanometer scale (16–19). Based on their tremendous structural diversity (20), plant lipids might account for the formation of plethora membrane domains (21).

The most complete and quantitative repertoire of the different molecular species of PM lipids and proteins is thus necessary to understand how they contribute together to build PM organization and dynamics. Different proteomes of purified plant PM have been characterized so far (for review see (22–24)) but the improvement in available genomic resources and mass spectrometry methods prompted us to reembark on such approaches. Besides, although the lipids present in the PM of plants have already been analyzed in different species (25,26), an overall picture reporting the precise amount and molecular description of each lipid species in this membrane is still missing, particularly in the model plant Arabidopsis acting as a reference for plant biologists. Building a reference of the whole lipidome and proteome of plant PM from a single material seems all the more necessary that variations of molecular compositions from one species, or one organ, to another, make it very difficult to get a global picture from data collected in different studies. So far, the only joint description of PM lipidome and proteome, has been performed on *Mesembryanthemum crystallinum* focusing on changes triggered by salt stress (27).

Here we report on the first joint extensive characterization of the Arabidopsis PM (further called AtPM) lipidome and proteome using state of art methods. Suspension cells were identified as a suitable starting material for several reasons: (i) they are cultivated in constant conditions (temperature, light, culture medium), and fully devoid of microbial contaminations, thus avoiding the risk of possible fluctuation in their composition triggered by biotic and abiotic stress (ii) their undifferentiated nature allows to avoid the compositional bias inherent in the study of differentiated tissues. The combined use of classical and newly developed methods allowed an in depth qualitative and quantitative characterization of the PM full lipidome and the use of sate of art proteomics methods and the benefit of the very recently actualized data bases yielded an unprecedented repertoire of PM proteins. The analysis of these two data sets in relation to each other gives a holistic view of the PM composition and provides clues to understand how the molecular characteristics of lipids and proteins are finely tuned to give a specific signature to the plant PM.

### Experimental procedures

#### Cell culture, PM preparation and purity

Arabidopsis cells ecotype Columbia-0 (Col-0) were grown in liquid culture consisting of Murashige and Skoog (MS) medium pH 5.6, containing MS salts, Nitsch vitamins, α-naphthalene acetic acid (0.5 mg/L), kinetin (0.05 mg/L) and sucrose (30g/L). Cell multiplication was carried-out by weakly dilution (20:100) into fresh medium under continuous light, 25 °C, and shaking (150 rpm). Microsome and PM preparations were performed at 4 °C. Microsomal proteins were extracted using the differential centrifugation procedure described by Stanislas et al. (28). Cells were collected by filtration during the exponential phase, frozen in liquid nitrogen, and homogenized with a Waring Blendor in grinding medium (50 mM Tris-MES, pH 8.0, 500 mM sucrose, 20 mM EDTA, 10 mM DTT, and 1 mM PMSF). The homogenate was centrifuged at 16,000 X *g* for 20 min. After centrifugation, supernatants were collected, filtered through two successive screens (63 and 38 µm), and centrifuged at 96,000 X *g* for 35 min. The resulting microsomal fraction was purified by partitioning in an aqueous two-phase system (polyethylene glycol 3350/dextran T-500, 6.6% each) to obtain the PM fraction (29). The sensitivity of ATPase activity to vanadate was used as PM marker (30). This protocol was repeated three times in order to obtain three independent microsomal (Atµ) preparations and three corresponding PM (AtPM) fractions. Protein and lipid profiling experiments were further performed on the latter PM fractions according to the workflow schematized in **Figure 1**.

**Figure 1.**
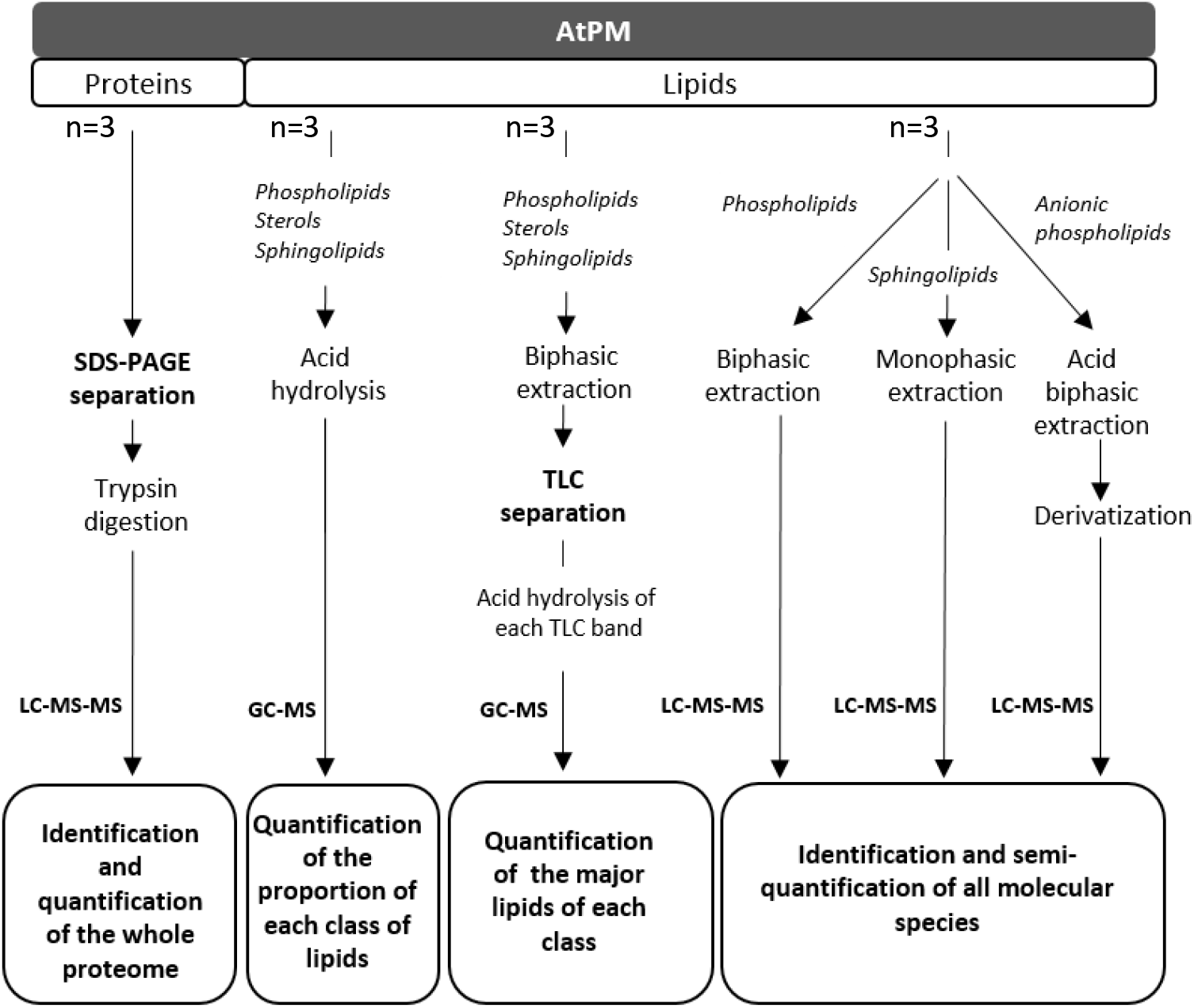
Rationale for the determination of AtPM proteome and AtPM lipidome. Plasma membrane purified form *Arabidopsis thaliana* suspension cells were split in two parts: one to identify and quantify the whole proteome, the other to identify and quantify the whole lipidome on three-fold methods. First, quantification of lipid moieties after hydrolysis and quantification by Gas Chromatography coupled to mass spectrometry (GC-MS) using internal standards. Second, quantification of each lipid class after separation by High-Performance-Thin Layer Chromatography (HP-TLC), scratched of the lipids, hydrolysis and further quantification by GC-MS) using internal standards. Third, identification and semi-quantification of all molecular species by Liquid-Chromatography coupled to MS using internal standards.

### Protein profiling

#### Sample pre-fractionation and protein digestion

For each biological replicate (n = 3), Atµ and AtPM proteins (10 µg) were pre-fractionated by a 0.5 cm migration on 12% SDS-PAGE. After Coomassie Brillant Blue staining, each gel was washed in distilled water, destained using 100 mM NH_4_CO_3_ in 50% acetonitrile (ACN), and then cut into 3-mm squares for subsequent proteomic analysis. A reduction step was performed by addition of 40 µl of 10 mM dithiotreitol in 50 mM NH_4_HCO_3_ for 30 min at 56 °C. The proteins were alkylated by adding 30 µl of 55 mM iodoacetamide in 50 mM NH_4_HCO_3_ and allowed to react in the dark at room temperature for 45 min. Gel pieces were washed in 50 mM NH_4_HCO_3_, then ACN, and finally dried for 30 min. In-gel digestion was subsequently performed for 7 h at 37 °C with 125 ng of modified trypsin (Promega) dissolved in 50mM NH_4_CO_3_. Peptides were first extracted successively with 0.5% (v/v) TFA, 50% (v/v) can, and then with pure ACN. Peptide extracts were dried and suspended in 25 µl of 0.1% (v/v) HCOOH, and 2% (v/v) ACN. Peptide extracts were dried and suspended in 25 µl of 0.1% (v/v) HCOOH, and 2% (v/v) ACN.

#### Liquid chromatography-tandem mass spectrometry (LC-MS/MS) analysis

Peptide separation was performed using a nanoelute UHPLC (Bruker, BillerikaUSA). Peptides were first desalted using a nanoEase Symmetry C18 TrapColumn, (100Å, 5 µm, 180 µm X 20 mm, Waters) with 0.1% HCOOH (v/v) in water for 1 min at 100 bar. Peptides were further separated on a Aurora C18 column (25 cm × 75 μm ID, 1.6 μm FSC C18, ionOptics). The mobile phase consisted of a gradient of solvents A: 0.1% HCOOH (v/v), in water and B: 100% ACN (v/v), 0.1% HCOOH (v/v) in water. Separation was set at a flow rate of 0.3 μl/min using a linear gradient of solvent B from 2 to15% in 60 min, followed by an increase to 25% in 30 min, 37% in 10 min, 90% in 5 min and finally to 90% for 10 min. Eluted peptides were analyzed with a timsTOF Pro (Bruker) using a Captive spray interface. Ionization (2 kV ionization potential) was performed with a liquid junction. Peptide ions were analyzed using Bruker otofControl 6.2.0 on PASEF Mode with tims activated, a m/z range from 100 to 1700, and a mobility ramp from 0.60 Vs/cm² to 1.6 Vs/cm². Each acquisition cycle was performed with 1 MS of 100ms, followed by 10 PASEF frames of 100 ms. For each PASEF frame, 10 MS/MS spectra were acquired with a rolling collision energy from 20eV to 59eV. Isolation was set at 2m/z for precursor mass less than 700 m/z and 3m/z for others. A dynamic exclusion was performed at 0.4min. Precursor was re-analyzed when intensity was 4 times higher than the previous scan to achieve a global MS/MS spectrum intensity of 20K counts.

#### Protein identification

Proteins were identified using X!Tandem (version 2015.04.01.1) (31) by matching peptides against the Araport11 database (www.araport.org) containing 48,358 protein entries downloaded from (https://www.araport.org/downloads/Araport11_Release_201606). Enzymatic cleavage was declared as a trypsin digestion with one possible missed cleavage in first pass. Cysteine carbamidomethylation were set to static, while methionine, protein N-terminal acetylation with or without excision of methionine, dehydration of N-terminal glutamic acid, deamination of N-terminal glutamine and N-terminal carbamidomethyl-cysteine as possible modifications. Precursor mass precision was set to 10 ppm with a fragment mass tolerance of 0.5 Da. Identified proteins were filtered and grouped using X!TandemPipeline (32) according to (i) the tolerated presence of at least two peptides with an *E-*value smaller than 0.01 and (ii) a protein *E-*value (calculated as the product of unique peptide *E-*values) smaller than 10^−5^. These criteria led to a false discovery rate (FDR) of 0.3% for peptide and protein identification.

#### Experimental design and statistical rationale

Protein abundance values were calculated by spectral counting using normalized spectral abundance factor (NSAF) analysis that is based on the cumulative sum of recorded peptide spectra matching to a given protein (33). This method estimates protein abundance by first dividing the spectral count for each protein by the protein length and then dividing this number by the sum of all length normalized spectral counts for each organism and multiplying by 100. A NSAF value was calculated for each protein across the six samples (three biological replicates for each microsomal and PM preparations). When necessary for statistical analysis, NSAF percentages were arcsine square root*-*transformed to obtain a distribution of values that could be checked for normality using the Kolmogorov-Smirnov test. Significant (*p*-value < 0.05) differences between transformed NSAF values were analyzed using the Welch-test (degrees of freedom = n−1), which is compatible with unequal variances between groups (34).

#### *In silico* protein characterization

Digital immunoblotting that refers to the MS-based quantification of proteins having an exclusive cellular component localization (35), was performed by combining experimentally know localization with NSAF spectral counting. Experimentally known localization of proteins, as inferred from HDA (High Throughput Direct Assay) and IDA (Inferred from Direct Assay) GO experimental evidence codes for cellular components (http://geneontology.org), was searched against Araport11 (https://www.arabidopsis.org/) (36) and SUBA5 (https://suba.live/) (37) built on the TAIR10 Arabidopsis proteome.

Alpha-helical trans-membrane (TM) regions and length were predicted according to TMHMM-2.0 (https://services.healthtech.dtu.dk/service.php?TMHMM-2.0). The presence of N-terminal signal sequences (SPs) that drive protein translocation across or integration to membranes, was predicted according to SignalP 5.0 (https://services.healthtech.dtu.dk/service.php?SignalP-5.0). The GPS-Lipid server (http://lipid.biocuckoo.org/webserver.php) was used as the lipid modification predictor for N-myristoylation, S-palmitoylation, and prenylation (S-farnesylation and/or S-geranylation. Glycosylphosphatidylinositol (GPI) anchor (i.e., glypiation) prediction was performed using NetGPI-1.1 (https://services.healthtech.dtu.dk/service.php?NetGPI-1.1). GRAVY values for protein sequences, defined by the sum of hydropathy values of all amino acids divided by the protein length, were retrieved using the GRAVY calculator (https://www.gravy-calculator.de/). Isoelectric points (pI) were retrieved from SUBA5. The function of the identified proteins was sought using the Mercator 3.6 web-based pipeline (https://plabipd.de/), which divides proteins into 35 hierarchical, non-redundant functional classes using MapMan bin codes (38). Arabidopsis Acyl-Lipid Metabolism Database (ARALIP: http://aralip.plantbiology.msu.edu/; (39), and Uniprot (https://www.uniprot.org/) were also used for annotation of lipid-related proteins.

### Lipid profiling

#### Absolute quantification of lipid moieties by GC-MS

For the analysis of total fatty acids by GC-MS, 200µg of AtPM were transmethylated at 110 °C overnight in methanol containing 5% (v/v) sulfuric acid and spiked with 10 µg of heptadecanoic acid (C17:0) and 10 µg of 2-hydroxy-tetradecanoic acid (h14:0) as internal standards. After cooling, 3 mL of NaCl (2.5%, w/v) was added, and the released fatty acyl chains were extracted in hexane. Extracts were washed with 3 mL of saline solution (200 mM NaCl and 200 mM Tris, pH 8), dried under a gentle stream of nitrogen, and dissolved in 150 mL of BSTFA -trimethylchlorosilane. Free hydroxyl groups were derivatized at 110 °C for 30min, surplus BSTFA-trimethylchlorosilane was evaporated under nitrogen, and samples were dissolved in hexane for analysis using GC-MS.

For the analysis of sterols moieties by GC-MS, 200µg of AtPM were transmethylated at 85 °C, for a shorter time to avoid degradation of sterols, i.e., 3h in methanol containing 1% (v/v) sulfuric acid spiked with 10µg of cholestanol, as internal standard. After cooling, 3 mL of NaCl (2.5%, w/v) was added, and the released sterol moieties were extracted in hexane. Extracts were washed with 3 mL of saline solution (200 mM NaCl and 200 mM Tris HCl, pH 8), dried under a gentle stream of nitrogen, and dissolved in 150 mL of BSTFA trimethylchlorosilane. Free hydroxyl groups were derivatized at 120 °C for 30min, surplus BSTFA-trimethylchlorosilane was evaporated under nitrogen, and samples were dissolved in hexane for analysis using GC-MS.

GC-MS was performed using an Agilent 7890A gas chromatograph coupled MS detector MSD 5975-EI (Agilent). An appropriate capillary column was used with helium carrier gas at 2 mL/min; injection was done in splitless mode; injector and mass spectrometry detector temperatures were set to 250 °C. Injection (1µl) was done in splitless mode; injector and mass spectrometry detector temperatures were set to 250 °C. Ionization energy was set at 70eV with m/z ranging from 40 to 700. For LCFA, HP-5MS capillary column (5% phenyl-methyl-siloxane, 30-m, 250-mm, and 0.25-mm film thickness; Agilent) was used. The oven temperature was held at 50 °C for 1 min, then programmed with a 25 °C/min ramp to 150 °C (2-min hold) and a 10 °C/min ramp to 320 °C (6-min hold). For VLCFA, DB-23 capillary column (60-m, 250-mm, and 0.25-mm film thickness; Agilent) was used and the oven temperature was held at 50 °C for 1 min, then programmed with a 25 °C/min ramp to 210 °C (1-min hold) and a 10° C/min ramp to 306 °C and a 65 ° C/min ramp to 320 °C (3-min hold). For sterols moieties, HP-5MS capillary column (5% phenyl-methyl-siloxane, 30-m, 250-mm, and 0.25-mm film thickness; Agilent) was used and the oven temperature was held at 200 °C for 1 min, then programmed with a 10 °C/min ramp to 305 °C (2.5-min hold) and a 15 °C/min ramp to 320 °C. Quantification of sterols moieties and fatty acid methyl esters was based on peak areas, which were derived from total ion current, using the respective internal standards.

#### Biphasic extraction for sterol and glycerolipids analysis

200 µg of PM were used for TLC coupled with GC-MS analysis and 20 µg for LC-MS. Samples were extracted with chloroform:methanol (2:1, v/v) at room temperature. Polar contaminants such as proteins or nucleic acid were removed by adding 1 volume of 2.5% NaCl solution. After phase separation, the lower organic phase, which contains lipids, was harvested and the solvent was evaporated under a gentle flow of N_2_ gas.

#### Monophasic extraction for sphingolipids

Sphingolipids were extracted from AtPM in the inferior phase of isopropanol/hexane/water (55:20:25, v/v) at 60 °C for 1h according to Toledo et al. (40). The extract was dried and resuspended in chloroform/methanol/water 30/60/8, v/v.

#### TLC Analysis of glycerolipids and and GlcCer

High-performance thin-layer chromatography (HP-TLC) plates were Silicagel 60 F254 (Merck, Rahway, NJ). Lipid extracts were chromatographed in chloroform:methanol:isopropanol:KCl 0,25%:methylacetate:triethylamine (30:10:25:7.8:24.3, v/v). Lipids were located under UV after staining with primuline in acetone/water 80/20. Lipid bands were recovered from HPTLC plates and submitted to the acid hydrolysis and the GC-MS analysis, as described below.

#### TLC analysis of sterols

Sterols were extracted following the biphasic extraction, as described. High-performance thin-layer chromatography (HP-TLC) plates were Silicagel 60 F254 (Merck, Rahway, NJ). Extracted lipids were chromatographed in chloroform:methanol (6:1, v/v). Lipids were located under UV after staining with primuline in acetone:water (80:20 v/v). Lipid bands were recovered from HP-TLC plates and submitted to the acid hydrolysis and the GC-MS analysis, as described below.

#### LC-MS/MS analysis of phospholipids

For the analysis of phospholipids by LC-MS/MS, lipid extracts were dissolved in 100 μL of solvent A (methanol/water 6/4 (v/v) + 10 mM ammonium formate +0.1%HCOOH) containing synthetic internal lipid standards (PE 17:0/17:0, PI 17:0/14:1 and PC 17:0/14:1 from Avanti Polar Lipids). LC-MS/MS (multiple reaction monitoring mode) analyses were performed with a model QTRAP 6500 (ABSciex) mass spectrometer coupled to a liquid chromatography system (1290 Infinity II, Agilent). Analyses were performed in the negative (PE, PG, PI) and positive (PC) modes with fast polarity switching (50 ms). Mass spectrometry instrument parameters were as follows: Turbo V source temperature (TEM) was set at 400 °C; curtain gas (CUR) was nitrogen set at 30 psi; the nebulizing gas (GS1) was nitrogen set at 40 psi; the drying gas (GS2) was nitrogen set at 50 psi and the ion spray voltage (IS) was set at - 4500 or +5500 V; the declustering potential was adjusted between -160 and -85 V or set at +35V. The collision gas was also nitrogen; collision energy varied from -48 to -62 eV and +37 eV on a compound-dependent basis. MRM Transition are presented in **Supplemental Table S1**. Reverse-phase separations were performed at 40 °C on a Supercolsil ABZ plus 2.1x100 mm column with 120-Å pore size and 3-μm particles (Supelco). The gradient elution program was as follows: 0 min, 20% B (isopropanol/methanol 8/2 (v/v) + 10mM ammonium formate + 0.1% HCOOH); 10 min, 50% B; 54 min, 85% B. The flow rate was set at 0.20 mL/min, and 5mL sample volumes were injected. The areas of LC peaks were determined using MultiQuant software (version 2.1; ABSciex) for relative phospholipid quantification.

#### LC-MS/MS analysis of sphingolipids

For the analysis of sphingolipids by LC-MS/MS, lipids extracts were then incubated 1h at 50 °C in 2 mL of methylamine solution (7ml methylamine 33% (w/v) in EtOH combined with 3mL of methylamine 40% (w/v) in water (Sigma Aldrich) in order to remove phospholipids. After incubation, methylamine solutions were dried at 40 °C under a stream of air. Finally, they were resuspended into 100 μL of THF/MeOH/H_2_O (40:20:40, v/v), 0.1% HCOOH containing synthetic internal lipid standards (Cer d18:1/h17:0, Cer d18:1/C17:0, GlcCer d18:1/C12:0) was added, thoroughly vortexed, incubated at 60 °C for 20min, sonicated 2min and transferred into LC vials.

LC-MS/MS (multiple reaction monitoring mode) analyses were performed with a model QTRAP 6500 (ABSciex) mass spectrometer coupled to a liquid chromatography system (1290 Infinity II, Agilent). Analyses were performed in the positive mode. Nitrogen was used for the curtain gas (set to 30), gas 1 (set to 30), and gas 2 (set to 10). Needle voltage was at +5500 V with needle heating at 400 °C; the declustering potential was adjusted between +10 and +40 V. The collision gas was also nitrogen; collision energy varied from +15 to +60 eV on a compound-dependent basis. MRM transition are presented in **Supplemental Table S1**. Reverse-phase separations were performed at 40 °C on a Supercolsil ABZ+, 100x2.1 mm column and 5µm particles (Supelco). The mobile phase consisted of a gradient of solvents A: THF/ACN/5 mM ammonium formate (3/2/5 v/v/v) with 0.1% HCOOH and B:THF/ACN/5 mM ammonium formate (7/2/1 v/v/v) with 0.1% HCOOH. The gradient elution program for Cer and GlcCer quantification was as follows: 0 to 1 min, 1% B; 40 min, 80% B; and 40 to 42, 80% B. The gradient elution program for GIPC quantification was as follows: 0 to 1 min, 15% B; 31 min, 45% B; 47.5 min, 70% B; and 47.5 to 49, 70% B. The flow rate was set at 0.2 mL/min, and 5mL sample volumes were injected. For quantification the areas of LC peaks were determined using MultiQuant software (version 3.0; Sciex).

#### LC-MS/MS analysis of anionic phospholipids

725 µL of a MeOH/CHCl_3_/1M HCl (2/1/0.1 v/v/v) solution and 150 µL water were added to the samples. Following the addition of the internal standard when appropriate and of 750 µL CHCl_3_ and 170 µL HCl 2M, the samples were vortexed and centrifuged (1500 X *g*, 5 min). The lower phase was washed with 708 µL of the upper phase of a mix of MeOH/CHCl_3_/0.01M HCl (1/2/0.75 v/v/v). Samples were vortexed and centrifuged (1500 X *g* / 3 min) and the washing was repeated once. Then, samples were kept overnight at -20 °C.

The organic phase was transferred to a new Eppendorf and the methylation reaction was carried out. For this purpose, 50 µL of TMS-diazomethane (2M in hexane) were added to each sample. After 10 min, the reaction was stopped by adding 6 µL of glacial acetic acid. 700 µL of the upper phase of a mix of MeOH/CHCl_3_/H_2_O (1/2/0.75 v/v/v) was added to each sample which was then vortexed and centrifuged (1500 x *g*, 3 min). The upper phase was removed and the washing step was repeated once. Finally, the lower organic phases were transferred to new Eppendorf. Following the addition of 100 µL MeOH/H2O (9:1 v/v), the samples were concentrated under a gentle flow of air until only a drop remained. 80 µL of methanol were added to the samples, which were submitted to ultrasounds for 1 minute, before adding 20 µL water, and be submitted to 1 more-minute ultrasound. The samples were finally transferred to HPLC vials for analysis.

Analysis of methylated anionic phospholipids (41) were performed using a liquid chromatography system (1290 Infinity II, Agilent) coupled to a QTRAP 6500 mass spectrometer (ABSciex). The chromatographic separation of anionic phospholipid species was performed on a reverse phase C18 column (SUPELCOSIL ABZ PLUS; 10 cm x 2.1 mm, 3µm, Merck) using methanol/water (3/2) as solvent A and isopropanol/methanol (4/1) as solvent B at a flow rate of 0.2 mL/min. All solvents are supplemented with 0.1% HCCOOH and 10 mM ammonium formate. 10 µL of samples were injected and the percentage of solvent B during the gradient elution was the following: 0–20 min, 45%; 40 min, 60%; 50 min, 80%. The column temperature was kept at 40 °C. Mass spectrometry analysis was performed in the positive ionization mode. Mass spectrometry data were treated using the MultiQuant software (ABSciex). Nitrogen was used for the curtain gas (set to 35), gas 1 (set to 40), and gas 2 (set to 40). Needle voltage was at +5500 V with needle heating at 350 °C; the declustering potential was +10 V. The collision gas was also nitrogen; was set between 26 to 45 eV according to the lipid classes. The dwell time was set to 5 ms. MRM transitions are presented in **Supplemental Table S1**.

## Results

The rationale to reveal the lipidome and proteome of Arabidopsis plasma membrane (AtPM) is shown in **Figure 1**. Three independent microsomal and PM fractions purified from Arabidopsis suspension cells were split in two parts: one to identify and quantify the whole PM proteome, the other to identify and quantify the whole PM lipidome on a three-fold method: first, quantification of lipid moieties, then quantification of each lipid class, and finally identification and semi-quantification of all molecular species.

### Protein profiling

PM fractions were isolated from Arabidopsis Col-0 cell suspensions by applying the conventional two-phase partition procedure (29) to a preparation of total membranes (microsomal fraction). The purity of the three independent AtPM preparations from microsomes (Atµ) was estimated first from enzymatic assays. A major part (85% ± 1, n = 3) of the total ATPase activity associated with the PM fraction was vanadate-sensitive, indicating that PM contamination by tonoplast and other membranes displaying ATPase activities (ER, mitochondria, plastids) was at most 15%. The protein composition of the three microsomal fractions and the corresponding PM fractions were further quantitatively analyzed by LC-MS/MS after trypsin digestion of the samples. Using at least two peptides for correct protein assignment and a probability of peptide misidentification inferior to 0.01, a total of 3,948 non-redundant plant proteins were overall identified in the six fractions originating from microsomes (n = 3) and PM (n = 3) preparations (**Supplemental Table S2**).

To achieve a posteriori validation of the enrichment in PM proteins and depletion of contaminants with a larger set of protein markers, we implemented a digital immunoblotting workflow according to Zhang and Peck (35). To favor experimentally known localization over the use of *in silico* algorithmic predictors for protein localization (42), an experimentally known subcellular localization was searched for each protein against SUBA5 and Araport11, according to HDA (High Throughput Direct Assay) and IDA (Inferred from Direct Assay) GO experimental evidence codes for cellular components **(Supplemental Table S3**).

As summarized in **Supplemental Table S4**, out of the 3,948 proteins identified, 3,745 (94.8%) was significantly enriched in PM-resident proteins with a reproducible purity of 86% (±1.4, n = 3), purified PM fraction was 2.2-fold higher than that in the microsomal fraction, an enrichment superior to what previously observed on the basis of SUBA predictions (35). Relative to microsomes, the amount of Golgi, mitochondrial and cytosolic proteins was significantly decreased by more than tenfold in the PM fraction (**Figure 2**). Extracellular proteins, recorded as the most abundant contamination, did not exceed 3.1% of the PM fraction in terms of spectral counts (**Figure 2**). Depletion of other contaminants ranged from 2.3 (plastid, cytoskeleton) to 3 (nucleus, endoplasmic reticulum). Overall, digital immunoblotting supports the enrichment in proteins assigned to the PM with a purity estimated to 86%, in full consistency with the enzymatic characterization.

**Figure 2.**
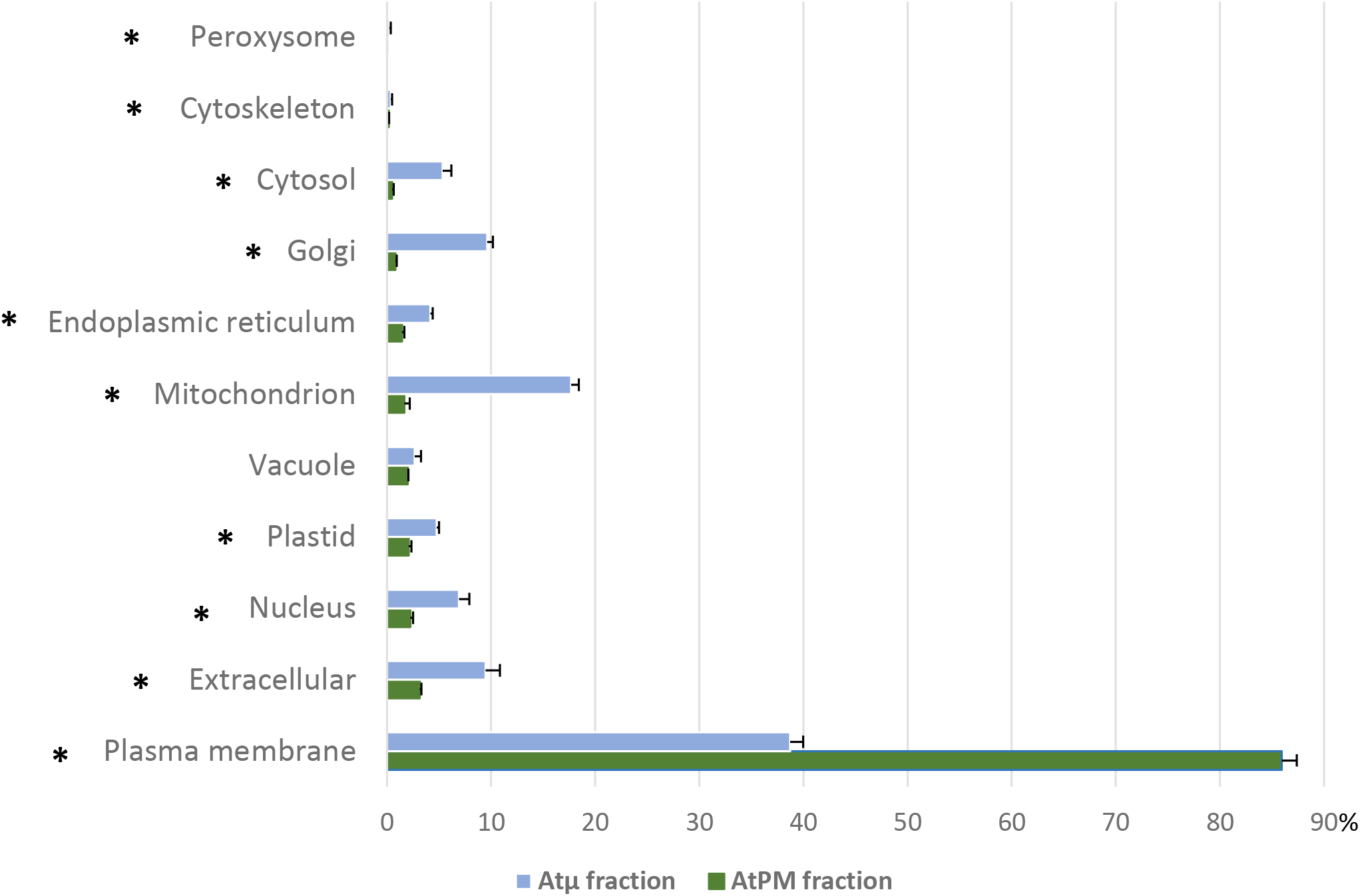
Comparative subcellular distribution of the proteins experimentally localized to an exclusive cellular component according to GO inference between microsomal (Atµ) and purified plasma membrane (AtPM) fractions. Histograms display the average cumulative sum (n = 3) of NSAF values (×100) for each cellular component. Bars represent the standard error of the mean. Asterisks refer to subcellular compartments displaying an abundance significantly (*p*-value < 0.05) different between the two fractions, as inferred from Welch-test analyses performed after angular transformation of cumulative NSAF data. Nota bene: the extracellular compartment includes cell wall proteins in SUBA5.

### An unprecedented repertoire of Arabidopsis PM proteins

Using the criteria described above for protein identification, a total of 2,994 proteins were identified in the three PM fractions, with 2,666, 2,281 and 2,875 proteins retrieved in the first, second and third AtPM preparation, respectively (**Supplemental Table S4)**. When retaining only those co-identified in the three replicates, the AtPM proteome of Arabidopsis Col-0 cells was hereby defined as a core-set of 2,165 proteins (**Supplemental Table S5**). Their abundance range spans four orders of magnitude and covers 98% of total PM spectral counts (**Supplemental Table S5)**. Among them, 194 proteins, which accounted for 2.3% of NSAF, were undetected in the three Atµ fractions (e.g., protein kinases, protein phosphatases 2C, oxysterol binding proteins, early nodulin-like protein), indicating that low abundant proteins have been enriched by the two-phase partition procedure. Many of the 2,165 identifications turned out to be canonical PM markers as determined by alternative methods for localization including fusion proteins with fluorescent reporters. Noteworthy, these PM landmarks included nitrate transporters (AT1G12110, AT4G21680), ammonium transporter 2 (AT2G38290), protein phosphatase 2C (AT3G51370), phospholipase D (AT4G35790), NOD26-like intrinsic protein (AT4G18910), YELLOW STRIPE like 2 protein (AT5G24380), cellulose synthase (AT5G09870), BRI1-like proteins (AT3G13380, AT1G55610), respiratory burst oxidase (AT1G64060), BAK1-interacting receptor-like kinase 1 (AT5G48380) and remorins (AT2G45820, AT3G61260) (**Supplemental Table S5**).

An experimentally known residence in the PM of Arabidopsis was searched for each of the 2,165 proteins against SUBA5 and Araport11. According to HDA and IDA evidence codes for cellular components, we retrieved 1,759 (81%) proteins previously recorded as PM proteins (Supplemental **Table S6**), resulting in the reproducible identification of 406 new PM-associated proteins, (**Supplemental Table S7**). Experimental evidence indicated that new AtPM proteins consisted of a small set (18.7%) of proteins exclusively localized to the PM with 81.3% of multi-localizing proteins (MLPs) (**Table S7**). The extracellular space and the Golgi apparatus were the two major co-localizing cellular components for the 112 MLPs, each accounting for 74 and 69 proteins, respectively.

### Physical and chemical characteristics of the Arabidopsis PM proteome

We examined the putative mechanisms by which the core-set of 2,165 proteins associate to the PM using web-based predictor tools (**Supplemental Table S5**). **Figure 3A** shows that the 736 proteins (34%) predicted as integral membrane proteins with one or more hydrophobic TM domains, accounted for 35% of total NSAF, as previously observed in Arabidopsis (43). A much higher proportion of NSAF abundance (65%) was assigned to proteins with no predicted TM domains (1,429 proteins). Among them, lipid-anchored proteins covalently attached to different FA acyl chains on the cytoplasmic side of the cell membrane via palmitoylation, myristoylation, or prenylation, represented the most abundant category (34% of spectral counts), with S-palmitoylation being the prominent single lipid modification targeting peripheral proteins to the membrane (NSAF of 16.6%). Overall, a total of 627 proteins (27% of PM spectral abundance) with no predicted TM domain displayed neither a SP nor a lipid anchor, among which a large majority (79%) was previously identified as PM resident proteins. These results point to alternative mechanisms mediating peripheral membrane association, such as lipid-binding domains (LBDs) that allow protein targeting to membranes by stereospecific interactions with a given phospholipid species (44,45). In Arabidopsis, 483 LBD-containing proteins have been described (46) and within the set of PM proteins lacking a TM domain and a lipid anchor, the identification for example of several C2-containing phospholipases D and Sec14-like proteins, support a role for LBDs in targeting proteins to the PM (**Table S5**). Because TM domain properties of integral membrane proteins act as major determinants of intracellular localization (47–49), we further assessed using the TMHMM-2.0 algorithm the quantitative distribution of TM domain-containing proteins in the AtPM proteome. Results indicated that single-spanning (i.e., bitopic) TM proteins ranked first in terms of abundance, accounting for 30% of the spectral counts recorded for all predicted integral membrane proteins (**Figure 3B**). Genome-wide bioinformatics analysis agree with this result, as they revealed that in Arabidopsis a large proportion (31%) of predicted bitopic proteins resides in PM (50). In addition, **Figure 3C** shows that the length of TM α-helical segments of bitopic proteins varied from 15 to 23 amino acids, with an overwhelming majority of 23-residue length, both in terms of protein number (77%) and cumulative protein spectral abundance (84%).

**Figure 3.**
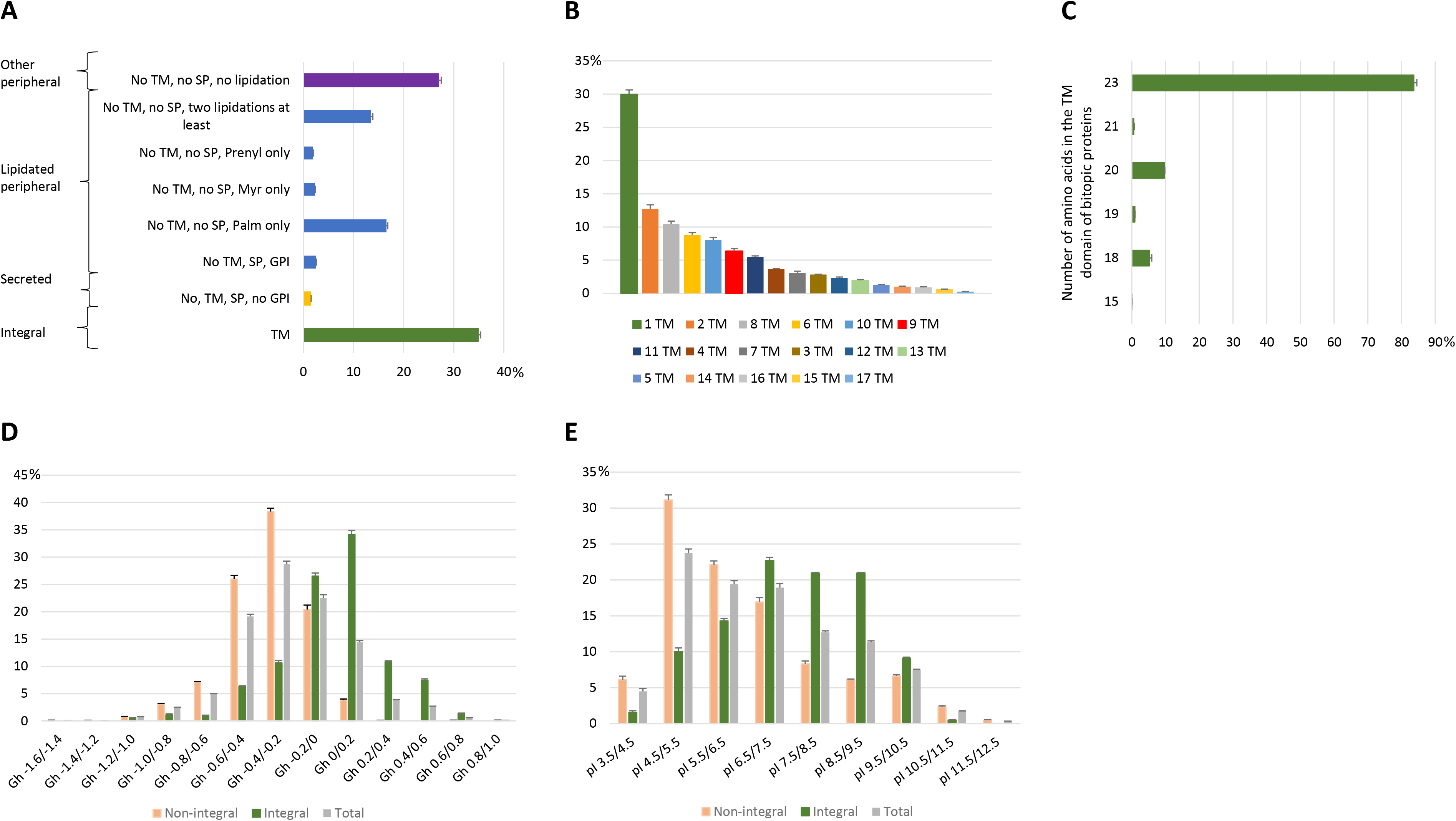
Physical and chemical characteristics of the Arabidopsis PM proteome. **A**, predicted membrane targeting mechanisms according to TMHMM-2.0, SignalP 5.0, NetGPI-1.1, and GPS-Lipid online tools, with TM, SP, Palm, Myr, Prenyl, and GPI refering to TM alpha-helix, N-terminal signal sequence, S-palmitoylation, N-myristoylation, prenylation, and glypiation, respectively. Distribution of **B**, single- and multi-spanning TM proteins, and **C**, TM segment lengths of bitopic proteins. Comparative distribution of **D**, GRAVY hydrophobicity and **E**, pI values between non-integral, integral and total proteins. Histograms display the average cumulative sum (n = 3) of NSAF values (×100) within each category. Bars represent the standard error of the mean.

The AtPM proteome was explored in terms of GRAVY score, with positive and negative values referring to hydrophobic and hydrophilic proteins, respectively. **Figure 3D** shows that the 2,165 proteins displayed a GRAVY index between -1.2 and +0.8, as observed in previous plant PM proteome repertoires (51,52), with 21% of NSAF corresponding to positive GRAVY scores. It also indicates a higher enrichment in hydrophobic proteins within the integral membrane protein fraction with 54% of spectral counts accounting for strictly positive GRAVY values relative to the peripheral fraction (4% of total NSAF), a result consistent with the hydrophobicity of TM segments (53). When examining the distribution of isoelectric points (pIs) retrieved in the AtPM proteome, **Figure 3E** indicates that 66.5 and 33.5% of PM spectral counts displayed a pI below and above 7.5, respectively. In agreement with this result, all proteomes analysed so far usually showed a bimodal pattern resulting in two sets of acidic and basic proteins (54). Nonetheless, pI distributions were quite different between predicted integral and non-integral AtPM proteomes with basic proteins accounting for 51.5 and 23.8% of spectral abundance, respectively. Of note, Schwartz et al. (55) similarly reported that integral membrane proteomes are enriched in basic proteins due to the basic residues commonly found on membrane spanning helices.

### Functional distribution of Arabidopsis PM proteins

To obtain a functional overview of the Arabidopsis PM proteome, we used Mercator for automated protein function annotation, according to the MapMan scheme. Among the 2,165 proteins co-identified in the three PM fractions, 1,818 proteins (84%) were assigned to 32 known functional categories. Two MapMan designations were missing, namely polyamine synthesis (numerical code 22) and micro RNA/natural antisense (numerical code 32), while the undefined/hypothetical function encompassed 347 proteins (**Table S5**). To assess their relative abundance, we added up each of the protein NSAF value recorded within a functional category. **Figure 4** shows the distribution of spectral counts within the eight MapMan categories displaying a NSAF sum superior to 3%. Signaling (e.g. MAP kinases, 14-3-3 proteins, G-proteins, receptor kinases), protein (e.g. synthesis, folding, modification, destination, degradation) and transport (e.g. ATPases, or transporters of sugars, amino acids, metabolites, ions, metals) were the three prominent functional group with a NSAF of 17, 16 and 15%, respectively. The next top-ranked functions included non-assigned proteins (9.3%), the cell category (8.7%) gathering proteins involved in cell organization, cell cycle, cell trafficking and vesicular transport, a pool of 8.7%, proteins responsive to stress, a family of miscellaneous enzyme (4.1%, e.g., glycosyl hydrolases, glutathione s-transferases, galactosidases), and proteins related to lipid metabolism (3.4%).

**Figure 4.**
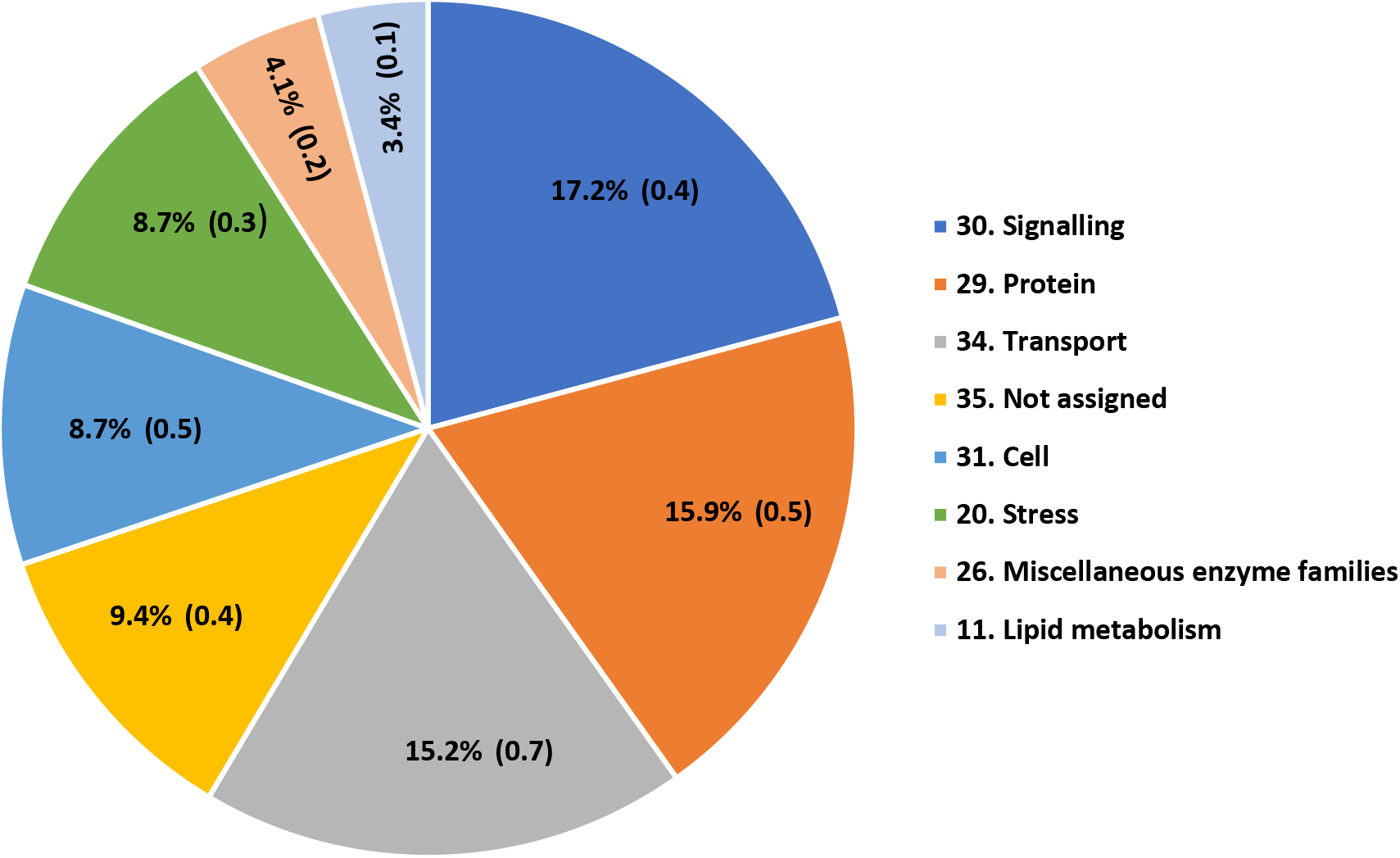
Functional distribution of the Arabidopsis PM proteome according to MapMan analysis. Only categories accounting for more than 3% of the total PM proteome abundance are presented. Results show the cumulative sum (n = 3) of NSAF values (×100) for each cellular component (mean ± standard deviation, n = 3).

To further draw a comprehensive map of lipid-related proteins in AtPM, we combined the lipid metabolism category (see above) with phosphoinositide signaling and lipid transport annotations using MapMan, ARALIP and Uniprot databases. Data mining overall retrieved a set of 133 lipid-related proteins (Supplemental **Table S8**), accounting for 6.1% of the 2,165 proteins co-identified in the three PM fractions. They included 18 proteins were involved in phosphoinositide signaling (e.g. PI4-kinases, PI-PLC, PI phosphatases), 29 in lipid transport (e.g. phospholipid-transporting ATPases, lipid transfer proteins), 36 in phospholipid metabolism (e.g. phospholipases, lysophospholipases, DAG kinases), 14 in sterol metabolism (e.g. sterol 3-beta-glucosyltransferase, delta(14)-sterol reductase), 6 in sphingolipid metabolism (e.g. LCB kinases) and 36 in FA metabolism (e.g. acyl-CoA oxidases, long chain acyl-CoA synthetases). Comparative NSAF abundance highlighted 62 proteins significantly (*p*-value < 0.05) enriched in AtPM versus microsomes (**Figure 5**). Four proteins (PI4K2, PLDBETA2, ORP1D and ORP2B) belonged to the set of new PM-associated proteins (**Figure 5**, **Table S8),** attesting for the depth of our study and its ability to evidence PM proteins of very low NSAF abundance. Of note, the PM enriched set encompassed most of the lipid-related proteins having role in lipid transport (24/29, 83%), phosphoinositide signaling (15/18, 83%), and phospholipid metabolism (17/36, 47%), as compared to the lower representation of sterol (4/14, 29%), sphingolipid (1/6, 16%), and FA metabolism (1/36, 3%).

**Figure 5.**
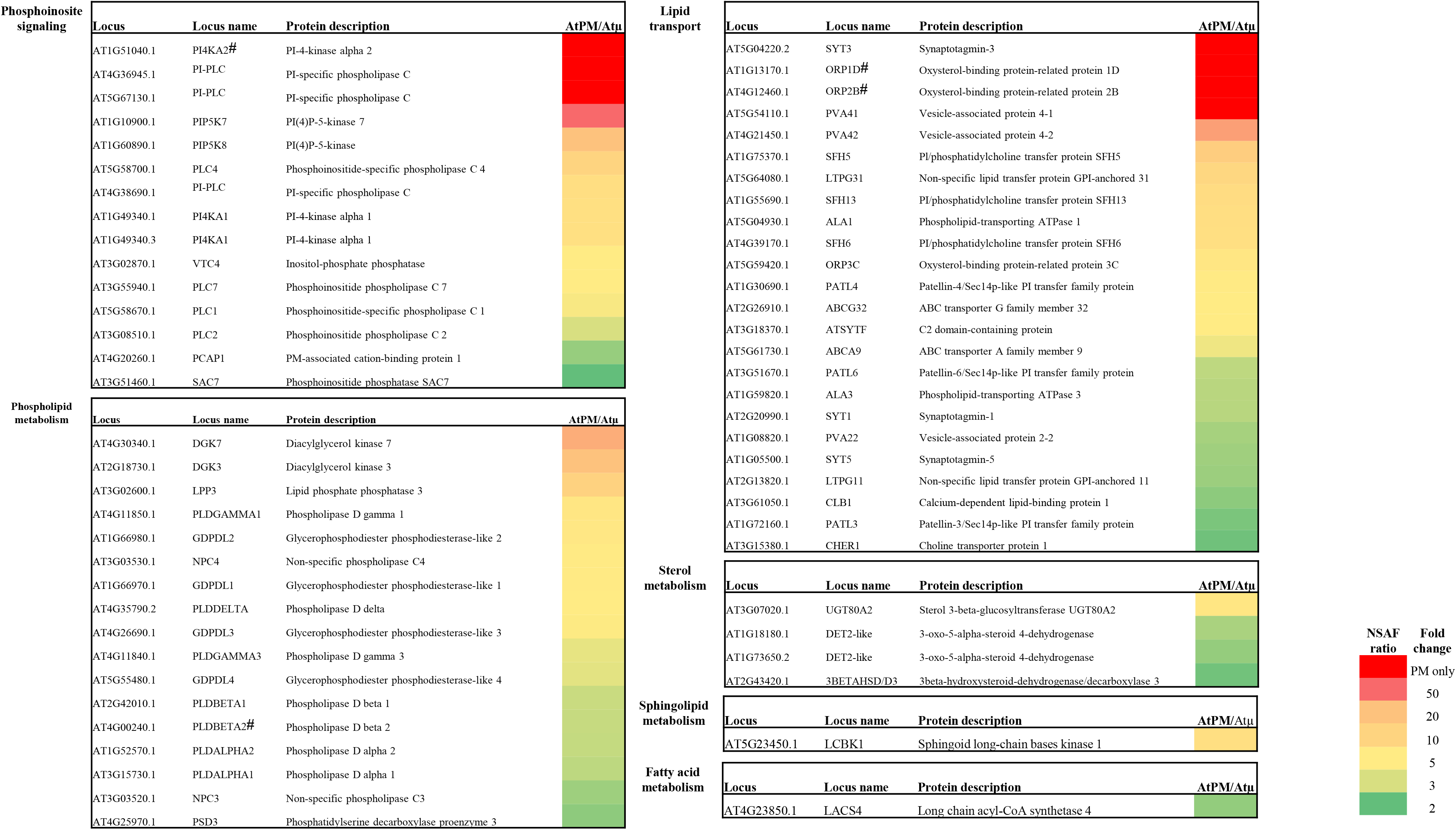
List of the 62 lipid-related proteins significantly enriched in in the Arabidopsis PM proteome relative to microsomes. For each protein, heatmap shows abundance fold-change between the plasma membrane (AtPM) and the microsomal (Atµ) fractions expressed as the mean (n = 3) NSAF ratio of AtPM to Atµ according to the color scale displayed on the right-hand side of the figure. The hash sign indicates proteins newly identified in the AtPM proteome.

### Lipid profiling

Lipidomic analysis have tremendously evolved in the last years. Mass spectrometry methods such as LC-MS/MS are widely used to determine the diversity of the molecular species. The quantification by LC-MS/MS is yet hampered because the ionization yield and the response of the MS analyzer depend on the nature of the lipid and the chemical environment, called the matrix. In addition, the molar response varies from one class of lipid to another as between homologs within a class depending on the nature of the fatty acid attached to the glycerol or to the long chain base backbone, and the energy collision (56). The need of appropriate standard for all lipids being a limitation, absolute quantifications using LC-MS/MS methods is very difficult. Therefore, following several previous works which tackle lipid quantification (57,58), we used here a combination of three different analytical methods for lipid identification and quantification (**Figure 1**). First, methanolic hydrolysis of PM lipids and analysis by gas chromatography coupled to mass spectrometry (GC-MS) allowed the quantification of sterols, long-chain fatty acids (FA), and very long-chain fatty acids (VLCFA) using internal dedicated standards. Second, the lipids were separated by high performance thin layer chromatography (HPTLC), scratched, hydrolysed and quantified by GC-MS. Third, the precise identification and regiolocalisation (when possible) of the molecular species of each lipid were determined by liquid chromatography coupled with mass spectrometry (LC-MS/MS) in multiple reaction monitoring (MRM) mode.

#### Quantification of the three classes of lipids

As described for the protein profiling, we first performed an absolute quantification of total lipids found in the microsomal and PM fractions, together with fresh Arabidopsis suspension cells. After full acid hydrolysis of lipids, corresponding lipid moieties were analyzed and quantified by GC-MS method (see Experimental Procedures). It was estimated that the amount of sterol moieties represents the total sterol content, and that LCFAs were mostly esterified in glycerolipids and (h)VLCFAs amidified in sphingolipids (59). Representative chromatograms are shown in **Supplemental Figure S1**. As expected, palmitic (C16:0) and oleic acid (C18:2) acids were the major LCFAs, even if chains harboring 18 carbon atoms with 0 to 3 double bounds were also detected (**Supplemental Figure S1A, B)**. VLCFA with up to 26 carbon atoms, odd and even number of carbons, 2-hydroxylated or not were also quantified, lignoceric acid (C24:0) and 2-hydroxylated with 24 carbon atoms acid (h24:0) being the major species (**Supplemental Figure S1A)**. Concerning the sterols (free and conjugated) analyzed after shorter acid hydrolysis, mostly β-sitosterol was detected in AtPM, with minor amount of campesterol but no stigmasterol (**Supplemental Figure S1C**).

The molar percentage of each class of lipids was estimated from these amounts of LCFA, (h)VLCFA and sterol moieties using an average molecular molar mass of 750g/mol for glycerolipids, 1200 g/mol for sphingolipids and 400 g/mol for sterols. **Figure 6A** shows that glycerolipids were by far the major lipids of fresh Arabidopsis suspension cells and microsomal fractions with only 10 mol% of sterol and 3 mol% of sphingolipids. These two latter classes were strongly enriched in the AtPM fraction, where glycerolipids, sphingolipids and sterols were distributed equally, with ca. 30 mol% each. Saturated FAs (mostly palmitic, lignoceric and h24:0) were strongly enriched in the AtPM compared to the microsomal fractions (**Figure 6B**), making the PM the most saturated membrane of the plant cell. The VLCFAs identified were mainly not hydroxylated and with FA chains up to C26 (**Figure 6C, D**). From AtPM lipid amount determination, the mass ratio of lipid/protein was calculated to be around 0.8 ± 0.03, in frame of previously described literature (59).

**Figure 6.**
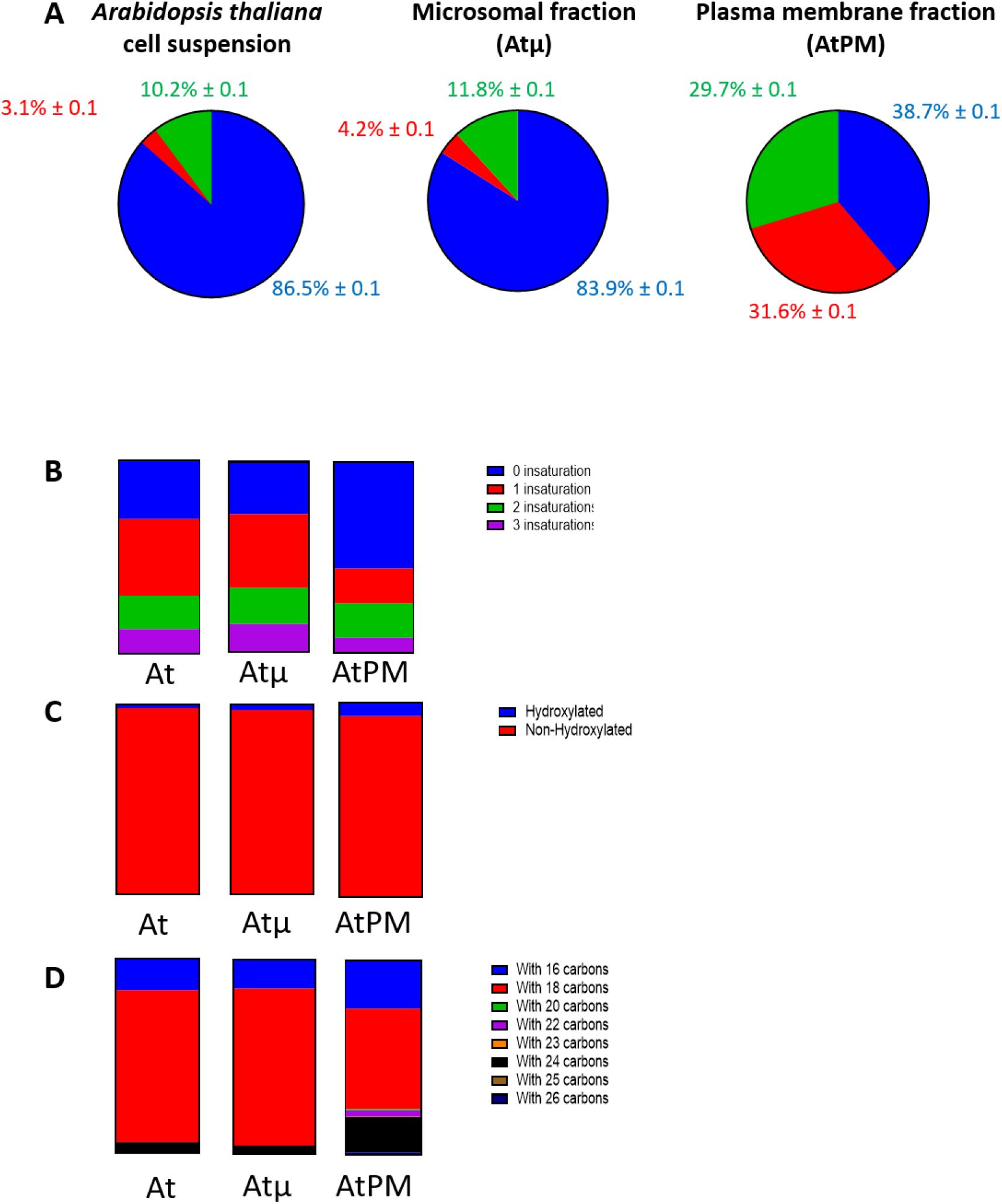
Quantification of the three lipid classes in fresh Arabidopsis suspension cell, microsomal fraction Atµ, and purified plasma membrane AtPM. **A**, Quantification by GC-MS of three major lipid classes namely sterols (green colour), glycerophospholipids (blue colour), and sphingolipids (orange colour) after acidic hydrolysis and derivatization. Experiments were done on fresh Arabidopsis suspension cells, microsomal (Atµ), and PM (AtPM) preparations. Results are shown as the molar percentage (mean ± standard deviation, n = 3). **B**, Unsaturation, **C**, Amount of non- and 2-hydroxylated fatty acids and **D**, number of carbon atom were calculated for the lipid quantified in fresh Arabidopsis suspension cells, Atµ and AtPM.

#### Quantification of the major lipids by TLC coupled to GC-MS

After separation by HP-TLC, individual lipid classes were scratched, hydrolyzed and quantified by GC-MS, allowing to determine the amount of AtPM major lipids. Note that minor lipids, barely detectable by TLC, were analyzed by LC-MS/MS (see below). First, HP-TLC allowed the separation of free and conjugated sterols. GC-MS quantification showed that free sterols were the most abundant followed by ASG and SG, with 83.7, 13.5, and 2.6 mol% of total sterols, respectively (**Figure 7A**). Second, glycerolipids and GlcCer were separated by HP-TLC. GC-MS quantification showed that PC and PE were the two major phosphoglycerolipids, representing 47.7 and 24.8 mol% of total glycerolipids, respectively, PI and PS representing 10.5 and 8.3 mol% of total phospholipids, respectively **(Figure 7B**). A few percent of PG were detected in AtPM whereas MGDG and DGDG were barely detectable (less than 0.1 mol%). Finally, GlcCer represents 16.8 mol% of total sphingolipids (**Figure 7C**). We failed to separate GIPC by HPTLC likely because of the presence of residual PEG-Dextran in the lipid extract perturbates the solvent migration. Therefore, we quantified them by LC-MS/MS. As plant GIPC standards are not commercially available, cauliflower GIPC, which are very similar to Arabidopsis GIPCs, were purified and quantified by GC-MS (**Supplemental Figure S3A, B)**. AtPM extracts were injected in LC-MS/MS using purified cauliflower GIPC as external standards. This allowed us to fully exploit the LC-MS/MS data and estimate the GIPC amount in AtPM. We determined that Hex(R1)-HexA-IPC and Hex-Hex(R1)-HexA-IPC represent ca. 63.8 and 10.3 mol% of total sphingolipids, respectively (**Figure 7C)**.

**Figure 7.**
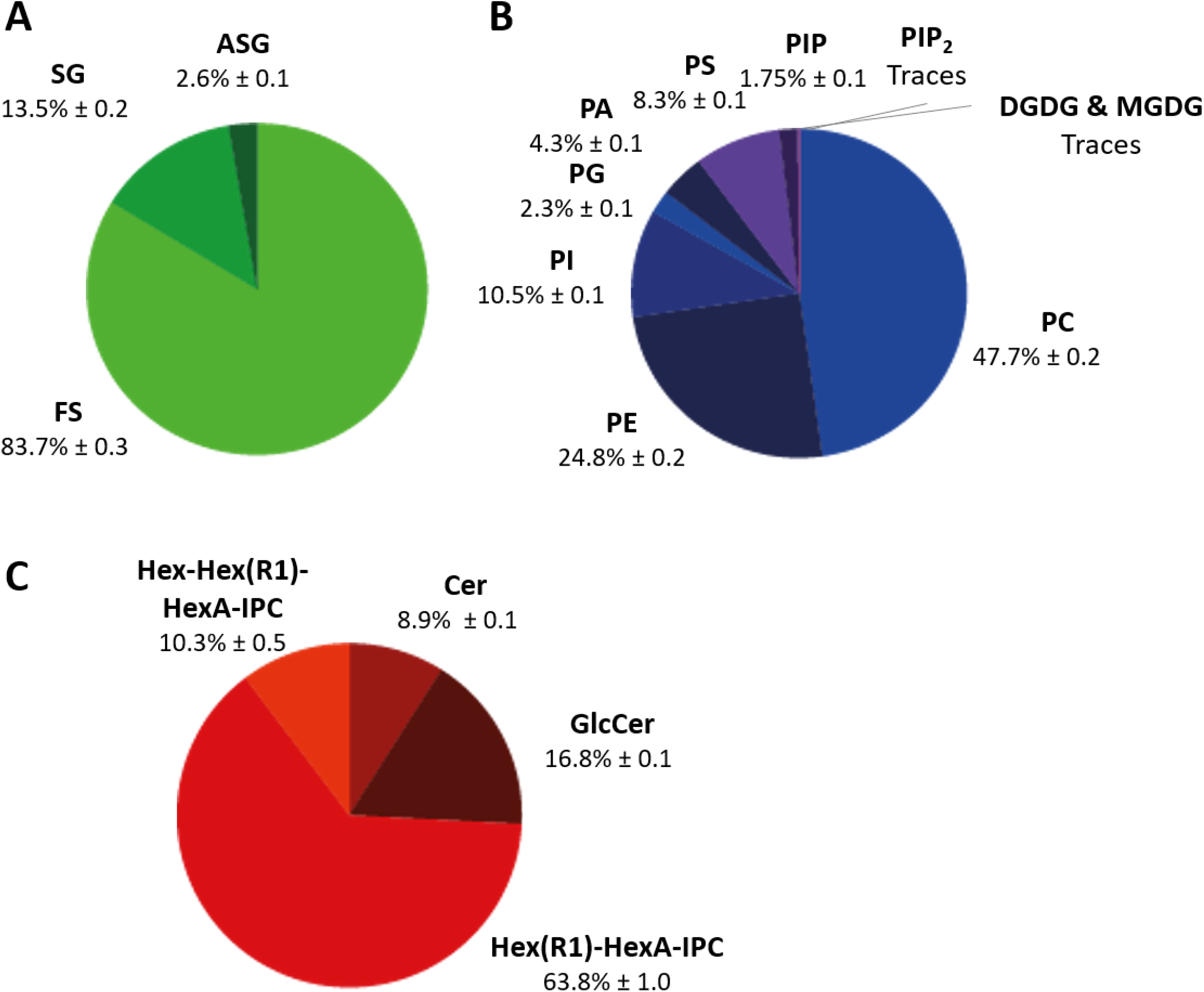
Lipidome of the Arabidopsis plasma membrane. **A**, Quantification by HPTLC coupled to GC-MS of sterol classes (FS, free sterol; SG, steryl glycoside; ASG, Acyl steryl glycoside). **B**, Quantification by HPTLC coupled to GC-MS of major glycerolipid classes (PC, PE, PI). Anionic lipids (PA, PS, PI, PIP, PIP_2_) were quantified by LC-MS/MS (see Figure 8), and quantification were integrated in the pie chart. **C**, Quantification by HPTLC coupled to GC-MS of GlcCer. Ceramides (Cer) and GIPC (Hex(R1)-HexA-IPC and Hex-Hex(R1)-HexA-IPC) were quantified by LC-MS/MS (see Figure 8), and quantification were integrated in the pie chart. Results are shown as the molar percentage (mean ± standard deviation, n = 3).

#### Extensive identification and regiolocalisation of lipid molecular species by LC-MS/MS

To obtain a precise determination of molecular species (fatty acid content and position), the different lipid extracts (see Experimental Procedures) were analyzed by LC-MS/MS. We set up MRM methods dedicated to the detection of major glycerolipids (PC, PE, PG, MDGD, DGDG), anionic phospholipids (PA, PS, PI, PIP, PIP_2_) and sphingolipids (LCB, LCB-P, Cer, GlcCer and GIPC series: Hex(R1)-HexA-IPC, Hex-Hex(R1)-HexA-IPC). These MRM methods can potentially detect 964 different molecules, see **Supplemental Table S1. Supplemental Figure S2** presents the results obtained for each class of lipids.

For glycerolipids, we showed that PC contain mostly C16 and C18 FA. The major PC species were 34:1, 34:2, 36:1, and 36:3. No VLCFA-containing PC were detected. PE molecules also contain C16 and C18 FA, with few percent of VLCFA up to 24 carbon atoms, with 34:1, 34:2 and 36:3 being the major species. Fragmentation of PE molecules allowed the regiolocalisation of FA found in *sn*-1 and *sn*-2. (Note that this was not possible for PC because it lost the polar head, and not the fatty acid in positive ionization mode). This revealed that major PE molecular species were PE 34.2 (16:0/18:2), PE 34.1 (16:0/18:1), and PE 36.3 (18:2/18:1). Traces of MGDG 34:2, 34:3 and DGDG 34:2, 34:3 were detected but represented less than 0.1% of total glycerolipids (**Figure 7B**, **Supplemental Figure S2**).

Sphingolipids were further analyzed by LC-MS/MS as described in Groux et al. (60). We failed to detect LCB- and LCB-phosphate, probably because they were below the limit of detection. The major ceramide species were t18:1/C24:0, t18:1/h24:0 and t18:1/h24:1. The major Hex-Hex(R1)-HexA-IPC were d18:0/h24:0 and d18:0/h24:1 (**Supplemental Figure S2**).

#### Analysis of anionic phospholipids

Anionic phospholipids like PA, PS, PI, PIP, and PIP_2_ were barely detectable by TLC or by LC-MS/MS following the classical method described previously. Thus, a dedicated derivatization step was added to increase the sensitivity of detection (**Figure 1)**, as described in Genva et al. (41), Ito et al. (61), and Yin et al. (62). This derivatization step by methylation using diazomethane remarkably enhanced the ionization, the separation of each peak and the limit of detection and quantification for each individual anionic phospholipid, more particularly PA, PS, and PIP_2_. Applied to AtPM, this sensitive method gave access to the determination of molecular species of anionic lipids.

Relative quantification using internal standards revealed that PI 34.1 and PI 36.1 represent the major PI molecular species (**Figure 8**). The major molecular species of PS contain VLCFA with 42:1, namely fatty acids up to 24 to 26 carbon atoms (**Figure 8**). PS represent 8.3 mol% of total glycerolipids (**Figure 7B**). We also determined that the major molecular species of PA were 34:1 and 36:1, whereas no VLCFA was detected in PA (**Figure 8**). PA represent around 4.3 mol% of total glycerolipids (**Figure 7B**). Concerning the polyphosphoinositides, PIP represent only 1.75 mol % of total phospholipids with 34:1 and 36:1 species. Only traces of PIP_2_ were detected with also consisting of 34:1 and 36:1 species (**Figure 7B, 8**).

**Figure 8.**
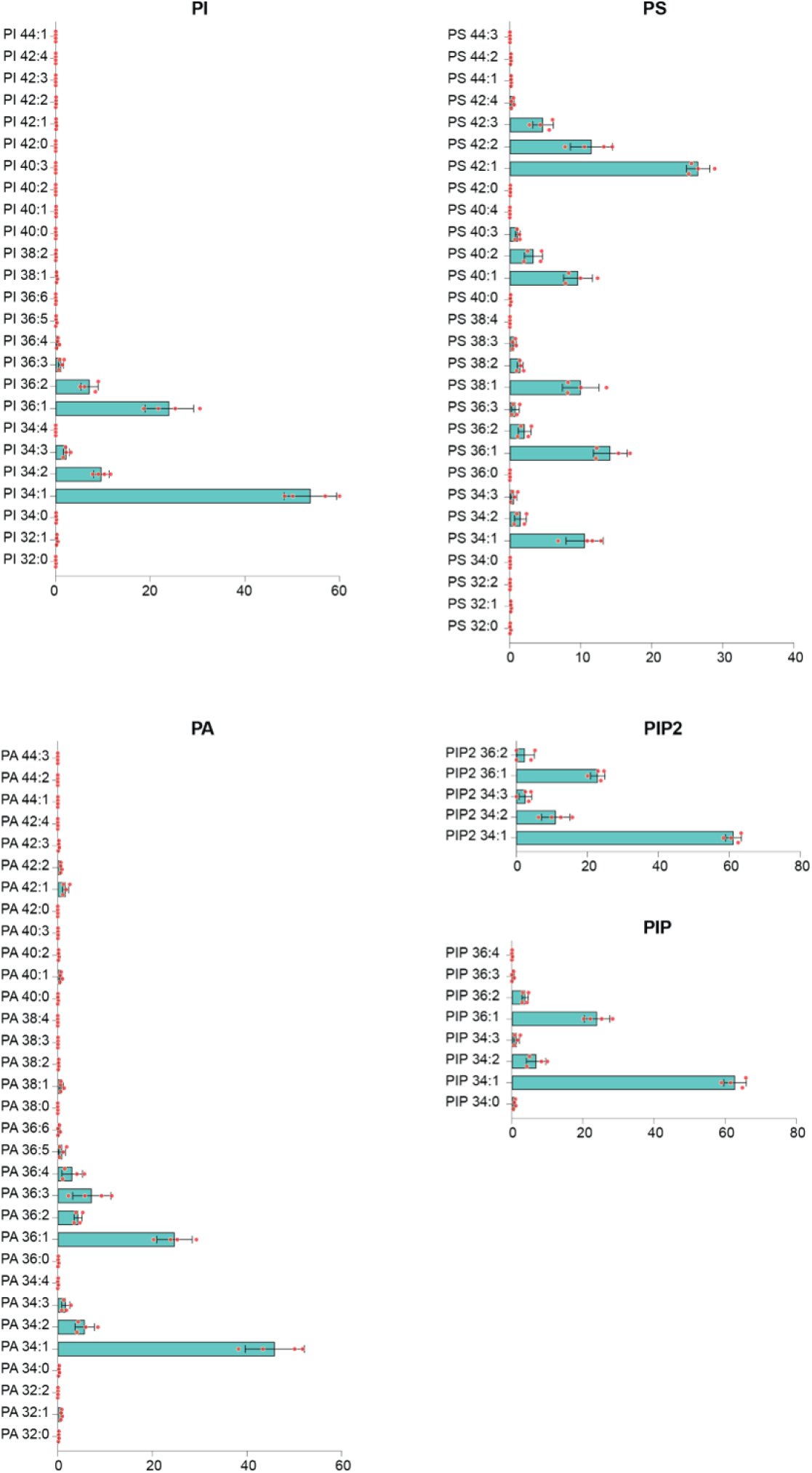
LC-MS/MS profiling of minor anionic lipid classes found in AtPM. AtPM anionic lipids namely PI, PS, PA, PIP, PIP_2_ were submitted to methylation by diazomethane and injected to LC-MS/MS. Results show the different molecular species expressed as molar percentage (mean ± standard deviation, n = 3).

### Full lipidome of AtPM

To summarize, we collected all the lipid profiling data with the major molecular species of each lipid and present them in a single figure, see **Figure 9**. Up to 405 different molecular species in AtPM were detected with GIPC (Hex(R1)-HexA-IPC d18:1/h24:0), PC 36:3, PE 34:2 (16:0/18:2), PS 42:1 and β-sitosterol being the 5 major lipids found in PM purified from *A. thaliana* suspension cells. Importantly, only traces of MGDG were detected confirming the lack of plastidial contamination, in line with the proteomic data (**Figure 2)**. Similarly, DGDG was also barely detectable. The weak amount of PA, less than 2 mol% (**Figure 9**), emphasized the lack of lipase activation (mostly phospholipase D) during AtPM purification and lipid extraction, a drawback frequently observed during plant membrane lipid analysis (63).

**Figure 9.**
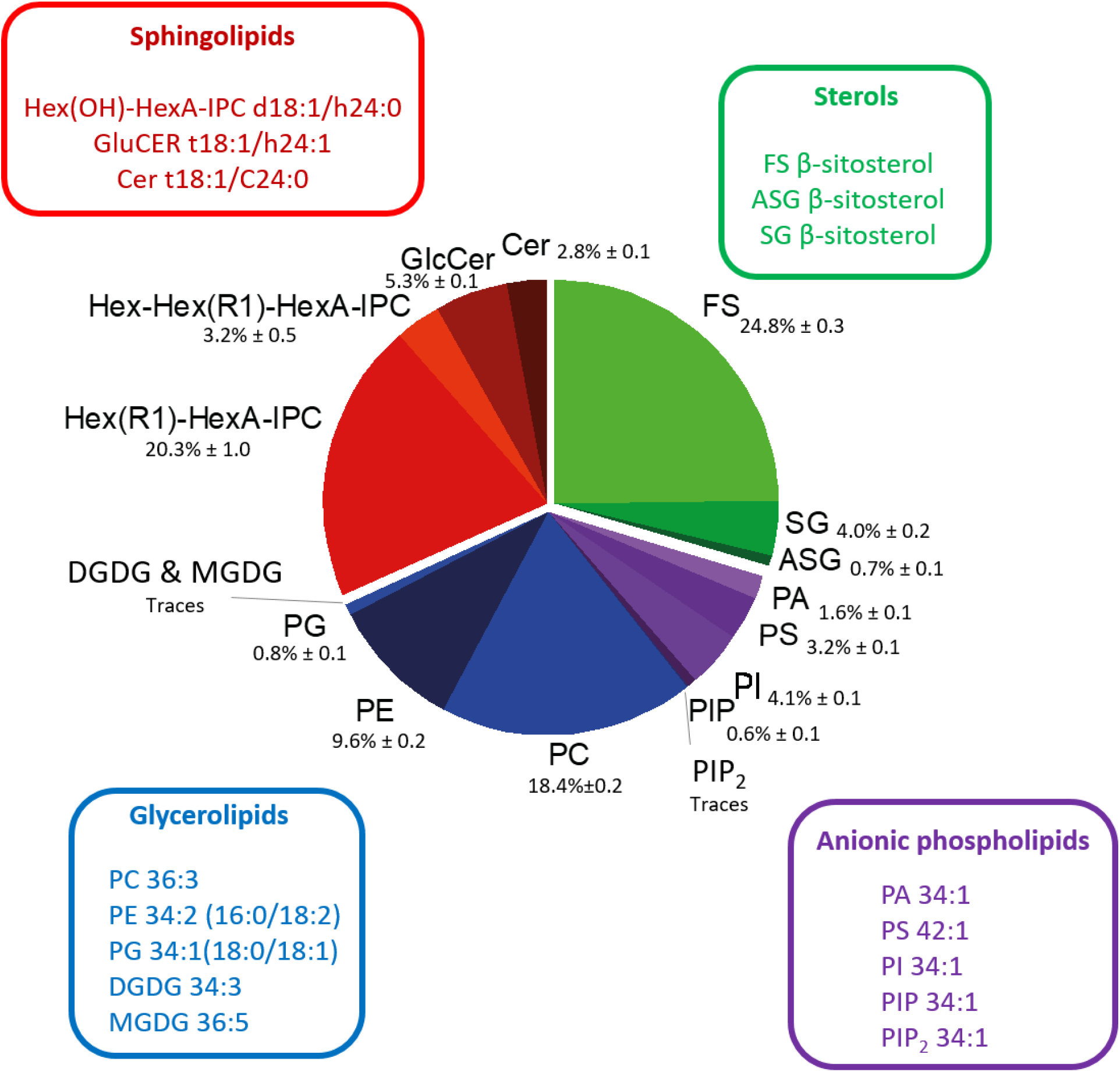
Lipidome of the Arabidopsis plasma membrane. Lipidomic results were gathered in a glucosylceramide, Cer; ceramide.

Concerning phosphoinositides, PIP represent 0.6 mol % of total lipids and only traces of PIP_2_ were detected (**Figures 7B, 8**). This is sound with previous work estimating PIPs to reach 0.2–1% of total cellular phospholipids, as reviewed in Balla et al. (64). Unfortunately, our LC-MS/MS method does not allow to characterize the phosphate isomers of these latter polyphosphoinositides, but previous work using genetically encoded biosensors specific to the polar head of phosphoinositides showed that PI4P is the major isomer found in plant PM (65,66).

The analysis of FA chain unsaturation of AtPM lipids clearly confirms the results of **Figure 6** with a high proportion of saturated or mono-unsaturated fatty acids present both in glycerolipids and sphingolipids, together with the presence of VLCFA in PS and sphingolipids (see **Supplemental Figure S4)**. PC and PE present a wider range of FA unsaturation than PS and PI.

## Discussion

### An increased quantitative coverage of the AtPM proteome

Numerous proteomic studies have been conducted to decipher the PM protein composition of Arabidopsis (35,43,51,67–72), but significant progress in the field of resource annotations (36,37) require an updated view of the AtPM proteome. With a total of 2,994 PM proteins identified (**Supplemental Table S2**), and 2,165 proteins reproducibly identified across three replications (**Supplemental Table S5**), the current analysis performed on cell suspensions has generated by far the most comprehensive qualitative and quantitative view yet for Arabidopsis, including 406 new PM-associated candidates. **Supplemental Table S9** provides a numerical comparison indicating the improvement in PM protein identifications over time, with a strong positive correlation (R^2^ = 0.98) between proteome coverage and available genomic resources (**Supplemental Figure S5**). Besides this linear relationship indicating that our characterization of AtPM benefited from the complete reannotation of the Arabidopsis reference genome (Araporort11, (36)), the identification of in the current PM fraction of many proteins previously indexed in TAIR and SUBA5 but undetected in microsomes, also argued for a better resolution of mass spectrometry analysis.

The accuracy of proteomic results after subcellular fractionation largely depends on the purity of the targeted compartment, as influenced by the degree of protein enrichment and extent of contamination from other subcellular fractions (22,23). In this work, we first monitored plasma membrane protein enrichment by the PM-specific ATPase activity, and further performed a posteriori validation through a digital immunoblotting based on the MS-based quantification of proteins experimentally known to have an exclusive cellular localization according to the recent resources Araport11 and SUBA5 (see **Figure 2**). Each approach independently substantiated a reproducible AtPM purity in the range of 85 to 86%, similarly to what observed in earlier studies (51,67).

The further *in silico* analyses we performed on the 2165 PM protein set fitted typical traits of the plant PM, as inferred from functional grouping, membrane-anchoring mechanisms, hydrophobicity pattern, pI distribution, and PM-characteristic features of bitopic proteins (**Figures 3, 4**). They also highlighted that the AtPM proteome mostly consisted of MLPs, as inferred from experimental evidence codes (**Tables S6, S7, S8**), arguing for proteins shuttling or operating across organelles to regulate metabolic reactions or to transmit information from the surrounding environment (73). The plant PM is notably known to form ER–PM membrane contact sites (MCSs), defined as tight junctions tethered by specific proteins (74,75). In support for a dual localisation of PM proteins to the ER, PVA11/VAP27-1, SYT3 (**Table S8**) and several multiple C2 lipid-binding domains and transmembrane region proteins (MCTP), namely AtMCTP3 (AT3G57880.1), AtMCTP4 (AT1G51570.1), AtMCTP6 (AT1G22610.1), AtMCTP15/QKY (AT1G74720.1), and AtMCTP16 (AT5G17980.1) were present in the AtPM proteome (**Table S5**). MCTPs that act as ER-PM tethers specifically at plasmodesmata, are proteins that insert into the ER via their transmembrane region while their C2 domains dock to the PM through interaction with anionic phospholipids (76). Likewise, SYT1 and SYT5 together with the Ca^2+^- and lipid-binding protein CLB1 form a complex enriched at ER–PM contact sites (77). Taken together, our results confirm that the proteomic approach succeeded in identifying and quantifying proteins closely associated to the PM.

### An exhaustive analysis yielding a complete picture of the lipid composition of Arabidopsis PM

Lipids represent a large and complex class of small molecules with a huge structural diversity which display hydrophobic and amphipathic chemical properties. The lipid composition of the PM has been studied in many plants like in corn (78), in the resurrection plant *Ramonda serbica* following dehydration and rehydration (79), in *A. thaliana* cell culture (80) or in *A. thaliana* plants with regard to cold acclimation (26), sterol biosynthetic mutants (70), sphingolipids composition (81), in oat and rye in relation to freezing tolerance (25), or more recently in the halophyte ice plant *M. crystallinum* (27). These works bring important piece of knowledge on the plant PM lipid content and its variation upon stress, but classes of lipids were often missing like GIPC or phosphoinositides because of lack of sensitivity of the methods (e.g., anionic lipids) or the use of not suitable lipid extraction procedures (e.g., GIPC).

Here, we decided to perform a very complete description of the full lipidome of the plant PM both qualitatively and quantitatively, with the determination of exact molecular species in terms of FA positions for some classes of lipids. Because of the complexity of lipids in terms of number of molecules and the lack of standards, our lipid profiling approach necessitates the use of various technics. LC-MS/MS is a very powerful method to decipher the molecular species of individual lipids, but the quantification is hampered by the lack of standards necessary to quantify the huge diversity of molecules because the detection is sensitive to the nature of the molecule especially with FAs. For example, among the 47,884 lipids species identified in the Lipid Maps database (https://www.lipidmaps.org), only 100 are commercially available as analytical standards. To obtain quantitative data, we thus added in our approach separation of lipids by TLC coupled to GC-MS as described in Khouri et al. (58), see **Figure 1**.

We built MRM tables that potentially allow the detection of almost a thousand different lipids molecular species (**Supplemental Table S1**). This enabled us to detect 405 different molecular species in AtPM, (**Figure 8**, **Supplemental Figure S2**), a number of molecules never reached in the study of the lipidome of the plant PM. Our study confirms previously published study on the lipidome of the plant PM, especially in *Arabidopsis* (80), but gave a further quantitative information (**Figure 9**). We thus showed that the three classes of lipids represent each 30 mol% and confirmed that GIPC namely Hex(R1)-HexA-IPC are the major sphingolipids in AtPM, with c24 and h24:0 fatty acids amidified on the LCB, as described in Grison et al. (80). This is also in line with the results obtained in PM purified from tobacco leaves and BY-2 suspension cells (59). GIPC were long forgotten in lipid analysis as they are not soluble in chloroform/methanol mixture and are hence lost in the interface or the aqueous phase. A suitable lipid extraction procedure (40) is required to ensure the full extraction of all classes of plant sphingolipids. In this work, we also measured the presence of more glycosylated Hex-Hex(R1)-HexA-IPC (series B) but 7-times less than series A GIPC. This is in agreement with *A. thaliana* as a conventional Eudicot plant in which GIPC with two saccharides are dominant (82). We have previously shown that both GIPC series are likely involved in the formation of membrane domains together with phytosterols (20,59). We note the presence of GIPC with di-hydroxylated LCB AtPM in suspension cells (**Figure S2**), a result at variance with previously reported GIPC in plant PM that contain mostly tri-hydroxylated LCBs (59,83). One can ask whether the LCB hydroxylase is down-regulated in suspension cells. We failed to detect LCB-LCB-phosphate, although these simple sphingolipids represent circa 1% of total sphingolipids (83). Either they are not present in the PM, or as amphiphilic molecules, they may have been lost during the process of PM purification.

Sitosterol and campesterol are the main phytosterols of AtPM, with the notable absence of stigmasterol in our model, Arabidopsis cell culture of ecotype Columbia-0 (Col-0). This appears at variance with Grison et al. (80) who showed that stigmasterol was the second most abundant phytosterols, using *A. thaliana* cell cultures of the ecoytpe Landsberg erecta. The activation/repression of the AtCYP710A1 gene that mediates stigmasterol production could be at play to explain this difference (84).

We also showed that PC and PE are the main phosphoglycerolipids with 24 or 26 carbon atoms with 0 to 4 insaturations, as described in Grison et al. (80). The absence of DGDG in our analysis showed that suspension cells growth medium was not deprived in phosphate (85). Finally, we developed a new protocol to measure the amount anionic lipids with PA, PS, PI and PIP representing 1.6, 3.2, 4.1 and 0.6 mol% of total lipids respectively, and only traces of PIP_2_ were detected (see **Figure 9)**. These results are in complete coherence with previously published work measuring the amount of lipids *in planta* using genetically encoded biosensors indicating that PIP are enriched in the PM, but not PIP_2_ (for review see (86)). LC-MS/MS also indicated that FA esterified in anionic lipids contain 16 or 18 carbon atoms with the notable exception of VLCFA found esterified in PS. These results are similar to what was published previously for the FA content of PA and PI in *A thaliana* (87). The DAG moieties of PIP and PIP_2_ are the same than the ones detected in PI, which indicates that polyphosphoinositides are metabolically derived from PI without any further fatty acyl editing. In contrast to PA, PI and PIPs, we found VLCFA esterified in PS (PS42:1, PS42:2 and to a lesser extend PS40:1 and PS42:3, see **Figure 8**). This is consistent with other previously published studies of PS extracted from Arabidopsis roots or seedlings analyzed by LC-MS/MS (65,87). The presence of VLFCA in PS seems a specificity of the plant kingdom. It is interesting to point that PS in animal cells mostly contains 18:0-20:4 fatty acids (88). The subcellular localization of PS varies during root cell differentiation and this gradient of localization tunes auxin signaling through the nano-clustering of the small GTPase ROP6 (89). To summarize, we provide here an LC-MS/MS method (41) that not only confirmed previous published results on the composition of individual classes of anionic phospholipids, but substantially improved their detection and quantification in one-shot.

### A lipid composition consistent with plant PM as an ordered mosaic of domains

Numerous evidences have pointed to the critical role of lipid acyl chains in membrane organization, in particular through two main properties: (i) the length of acyl chains, (ii) the ratio between saturated and unsaturated bonds along them (90), together with the presence of a hydroxylation at the 2 positions able to modulate the propensity to interact with sterols and other lipids (91). Using the quantitative data related to the 405 lipid molecular species here identified, global characteristics of Arabidopsis PM could be evidenced regarding such properties.

Noticeably, we found a clear predominance (56%) of fully saturated fatty acyl chains, the second in terms of representativeness being acyl chains harboring one unsaturation (18%) (**Figure 6**). A gradual enrichment of saturated PL species at the expense of monounsaturated species along the secretory pathway (ER > Golgi > PM) has been consistently evidenced in mammalian cells (92). This feature can be put in perspective with the increase in membrane order observed along the secretory pathway (88). Indeed, experimental data together with molecular dynamics simulation of varying membrane compositions showed that the proportion of saturated acyl chains is positively correlated with the order level of the membrane (92,93). In particular, vesicles formed from a total extract of plant PM lipids exhibit a slightly higher order level than vesicles formed of an equimolar mix of DOPC/DPPC/cholesterol (21). The high proportion of saturated FA chains together with sterols (**Figure 6**, see below) is thus in agreement with previous findings suggesting that the plant PM is highly ordered (20). The fundamental role of the monounsaturated/saturated ratio for elementary functions (from protective barrier to high trafficking) has been underlined and their biophysical characteristics mainly studied, either in a comparative way or in combination: namely, polyunsaturated PLs notably exhibit specific molecular properties, including a high capacity of adaptation towards mechanical stress (90,92). This agrees with the fact that PM from animal cells appear as “built for stability” whereas ER or Golgi membranes are characterized as highly dynamic, with significantly lower proportion of sterols and sphingolipids (94).

It is also interesting to consider the respective contributions of the different lipids to this global feature (see **Supplemental Figure S4**). It is clear than sphingolipids (GIPC and GlcCer) are mainly involved in the saturation of PM since more than 70% of their fatty acyl chains exhibit no unsaturation and the remaining 30% only one unsaturation. Regarding phospholipids, PS and PI are the more saturated with 75 and 80%, respectively, of their molecular species with one acyl chain saturated and the other monounsaturated. Conversely, for PC and PE molecular species with one acyl chain saturated and the other monounsaturated are a minority (about 30% each), whereas molecules with two or more (up to 6) unsaturations are prevalent. Results presented here thus point to a differential involvement of plant lipids in the various aspects of PM functioning, the “barrier function” being mainly ensured by sphingolipids (which also play a role in perception of environmental changes and subsequent signaling), whereas PC and PE might be mostly involved in the global adaptive processes. The presence of mainly monounsaturated PS is also consistent with the visualization of stable PS nanodomains on Arabidopsis root tip cell PM (89).

Another parameter to consider is the high diversity of molecular species present on the membrane, belonging to the different classes previously identified as major actors of membrane organization. Namely, it has been extensively described that the combination of lipids harboring specific properties due to their chemical structures is a key regulator of membrane organization, from the nano to the microscale (95). A noticeable information presented here is the presence on the PM of 30 mol% of sterols mainly sitosterol and campesterol which are the two free phytosterols with the highest ordering properties (21,96). Moreover, conjugated sterols (SG and ASG) which were also demonstrated as very efficient to increase membrane order, in combination with free sterols (21), have also been identified in significant proportion (13% and 3% of the total sterol amount, respectively). This contributes, together with saturation, to build the vision of a rather ordered plant PM. Moreover, sterols have been demonstrated to be not only key determinants of the plant PM global order *in vivo*, but also involved in the presence of ordered domains within this membrane (15,19–21,97). Sitosterol and campesterol have been proven particularly efficient in this respect, and the relative proportion of phospholipids (saturated and unsaturated), GIPC and sterols (free and conjugate) identified in this study is quite similar to the one providing a high level of ordered domain formation in the membrane of lipid unilamellar vesicles (21). This high amount of sterols, sphingolipids and saturated FA is thus fully supporting the presence within plant PM of ordered domains, in line with the « lipid raft » hypothesis that proposed rafts as PM nanodomains of high molecular order, enriched in cholesterol and sphingolipids, in which proteins involved in signaling can selectively interact with effector molecules (11). The fact that the VLCFAs of the sphingolipids (either GIPC or GlcCer) are mostly saturated is quite consistent with such a view of the plant PM.

### A finely tuned adjustment of the molecular characteristics of lipids and proteins

Global analysis of the molecular characteristics of PM proteins indicates than more than 80% of bitopic proteins (which represent by far the most abundant category of TM proteins identified) exhibit a transmembrane helix comprising 23 amino acids (**Figure 4**). These data support a model wherein long TM domains (≥ 23 amino acids) direct plasma membrane localization (98). This was further confirmed by Brandizzi et al. (99) who demonstrated that in plants among the different TM domains tested (from 17 to 23 aa), only TM segments with 23 aa drove proteins to the PM, while shorter TM domains (17 and 20 residues) remain mostly in ER and Golgi compartments, respectively.

The need to adjust the length of protein TM helices and the thickness of the hydrophobic core of the lipid bilayer to prevent or minimize a mismatch leading to a distortion of the bilayer or the protein, was identified for a long time as a crucial parameter (9). This suggested that the length of acyl chains, determining the thickness of the bilayer might be well suited to the length of protein domains spanning the membrane. This was very nicely demonstrated, comparing two species of the yeast *Schizosaccharomyces* with lipids exhibiting significant differences regarding the length of their FA chains: experiments coupled with bioinformatic analysis revealed that the length of protein TM domains were higher in the strain with the longest FA chains, and that reducing the carbon number of these FA chains resulted in a mistargeting of proteins (97). In the present study, lipidomic data indicate that about 30% of the lipids (phospholipids and sphingolipids) have one FA chains with at least 20 carbons. Thus, the characteristics of lipid FA chains and TM domains of bitopic proteins, specific to the PM, are very well adjusted one to another.

### The plasma membrane is enriched in proteins involved in phosphoinositide signaling, lipid metabolism and transport

Building on spectral counting and data mining of MapMan, ARALIP and Uniprot annotations, we here obtained a PM proteomic signature that functionally complements lipidomic insights (**Figure 5**). Indeed, among the 133 lipid-related proteins identified in the current study, most of the proteins associated to phosphoinositide signaling are enriched in the PM fraction compared to microsomes (**Figure 5**), three of them (PI4 kinase alpha2, PI-PLCs) being detected only in the PM fraction. This result is fully consistent with the recent findings indicating that PI4P, which mainly accumulates at the PM in plant cells (much more abundant than PIP_2_), support in plants many functions that are typically attributed to PIP_2_ in animal and yeast cells and should be considered a hallmark lipid of plant PM (96). Consistently, PI4-kinases alpha 1 and 2 involved in PI4P synthesis, are enriched in AtPM. These enzymes are tethered to the PM by a complex composed of proteins from the NO-POLLEN-GERMINATION EFR3-OF-PLANTS and HYCCIN-CONTAINING families (100), which here belong to the AtPM proteome (i.e., AT5G64090, AT2G41830, AT5G64090, respectively).

With regard to phospholipid metabolism, two isoforms of DAG kinases (DGK3 and DAGK7) were enriched in the PM fraction: these enzymes are involved in the production of PA in synergy with PI-PLC, an enzyme detected only in the PM in our analysis (**Table S**8). PI-PLC hydrolyses PIP_2_ to generate DAG, further phosphorylated by DGK to produce PA. The PA can also be generated through the hydrolysis of structural lipids (such as PC) by PLD: numerous isoforms of PLD are found enriched in the PM fraction (**Table S8**). The presence of this set of enzymes on the PM agrees with the very low amount of PIP_2_ and, conversely, the presence of a significant proportion of PA, which is a central lipid to mediate signaling processes in plants (101). The fatty acyl chains associated to PA (36:1 and 34:1) are also the most abundant in PIP_2_ as in structural lipids such as PC, in agreement with the involvement of these two pathways.

Related to sterol metabolism, we point to the PM enrichment and high abundancy of the sterol 3-beta-glucosyltransferase UGT80A2 (see **Supplemental Table S8**). This enzyme catalyzes the synthesis of steryl glycosides (SGs) and acyl steryl glycosides (ASGs). This is congruent with the identification of significant amounts of such conjugated sterols on the PM (respectively 13.5 and 2.6% of total sterols for SG and ASG, see **Figure 7**), and consistent with the fact that such enzymes can act on sitosterol, and campesterol which are the free sterols identified in this study.

Because PM does not participate in autonomous biogenesis of its structural lipids, they must be trafficked to their PM destination (102). This implies a constant flux of lipids from the synthesis compartments, mainly the ER, towards the PM, as mediated by vesicular and non-vesicular routes (103). Here, proteins having role in lipid trafficking were enriched for their large majority in AtPM relative to microsomes, including SYTs, VAPs, ORPs, and phosphatidylinositol transfer proteins (PIPT) of the SEC14-like family (**Supplemental Table S8**). SYT proteins are composed of a lipid transfer domain involved in lipid traffic between the ER and the PM (104). Lack of SYT1 and SYT3 increases the accumulation of DAG during cold stress, indicating an important role of SYTs in lipid metabolism and PM integrity (105). Of note, PI4P is critical for the localization of tethering proteins to the PM, including SYT1 (105) and MTCPs (76). VAP proteins interact with several lipid transfer proteins (LTPs), especially from the oxysterol-binding protein (ORP) family at membrane contact sites (106,107). Several ORPs transport their lipid ligand from the ER in exchange for PI4P on apposed membranes (108). PIPTs, which regulate a signaling interface between lipid metabolism and protein transport from the Golgi to the cell surface (109), include PM resident AtSFHs and AtPATLs, potentially involved in the regulation of the level and localization of phosphoinositides (110). Overall, these results point to the phosphoinositide lipid composition of the PM as a key determinant of membrane trafficking (65).

## Conclusions

This work provides the most complete repertoire to date of the lipids and proteins that make up plant cell PM. The functional and structural characteristics of the identified molecular species support elements already obtained by targeted studies and provide new insights into their ability to interact to shape the identity of this membrane.

This study clearly paves the way towards a crucial and still very poorly known characteristics: the asymmetry of composition and organization of the two leaflets of plant PM. Asymmetry has been thoroughly determined recently in erythrocytes but only few papers tackle this question in plants (111,112). Importantly, the description of the exact lipidome and the asymmetry is a pre-requirement to embark in the molecular dynamic modelisation of the plant PM, as it was reported in animal PM (113), and recently described in plant PM (114).

## Abbreviations

ACN: (acetonitrile)
ARALIP: (Arabidopsis acyl-lipid metabolism database)
ASG: (acylated steryl glycoside)
Atµ: (Arabidopsis microsomes)
AtPM: (Aradidopsis plasma membrane)
BSTFA: (*N*, *O*-bis(trimethylsilyl)trifluoroacetamide))
Cer: (ceramide)
Col-0: (columbia-0)
DAG: (diacylglycerol)
DGDG: (digalactosyldiacylglycerol)
ER: (endoplasmic reticulum)
FA: (fatty acid)
FDR: (false discovery rate),
G protein: (guanine nucleotide-binding protein)
GIPC: (glycosyl inositol phosphoryl ceramide)
GlcCer: (glucosylceramide)
GO: (gene ontology)
GPI: (glycosylphosphatidylinositol)
GRAVY: (grand average of hydropathicity index)
HDA: (high throughput direct assay)
HP-TLC: (high-performance thin layer chromatography)
hVLCFA: (2-hydroxylated very long chain fatty acid)
ID: (internal diameter)
IDA: (inferred from direct assay)
LBD: (lipid-binding domain)
LCB: (long-chain base)
LCB-P: (long-chain base-phosphate)
LC-MS/MS: (liquid chromatography-tandem mass spectrometry)
LTP: (lipid-transfer protein)
MCS: (membrane contact site)
MGDG: (monogalactosyldiacylglycerol)
MLP: (multi-localizing protein)
MS: (Murashige and Skoog)
MRM: (multiple reaction monitoring)
NSAF: (normalized spectral abundance factor)
PA: (phosphatidic acid)
PASEF: (parallel accumulation serial fragmentation)
PC: (phosphatidylcholine
PE: (phosphatidylethanolamine)
PG: (phosphatidylglycerol)
PI: (phosphatidylinositol)
PIP: (phosphatidylinositol monophosphate)
PI4P: (phosphatidylinositol 4-phosphate)
PIPT: (phosphatidylinositol transfer protein)
PIP_2_: (phosphatidylinositol bisphosphate)
PM: (plasma membrane),
PS: (phosphatidylserine)
PSI: (pound-force per square inch)
SG: (sterylglycosides)
SP: (N-terminal signal peptide)
SUBA: (subcellular localization database for Arabidopsis proteins)
TIMS: (trapped ion mobility spectrometry)
TM: (trans-membrane)
VLCFA: (very long chain fatty acid)

## Data availability

The mass spectrometry proteomic data have been deposited to the ProteomeXchange Consortium via the PRIDE (https://www.ebi.ac.uk/pride/) partner repository with the dataset identifier PXD040971. (Username; Password: kDUq3COy).

The mass spectrometry lipidomic data have been deposited in the public repository Zenodo (https://zenodo.org) under doi.org/10.5281/zenodo.7925332.

## Supplemental data

This article contains supplemental data including figures.

## Acknowledgments

The authors wish to thank Jérôme Fromentin for his expertise and support in the generation and maintenance of cell cultures, and Arnaud Mounier for the bioinormatic analysis of proteomic data. We thank Bordeaux-Metabolome platform for lipid analysis (https://www.biomemb.cnrs.fr/en/lipidomic-plateform/) supported by Bordeaux Metabolome Facility-MetaboHUB (grant no. ANR–11–INBS– 0010 to LF, SM). Proteomics analyses were performed on the PAPPSO platform (http://pappso.inrae.fr) which is supported by INRAE (http://www.inrae.fr), the Ile-de-France regional council (https://www.iledefrance.fr/education-recherche), IBiSA (https://www.ibisa.net) and CNRS (http://www.cnrs.fr). This study received financial support from the French government in the framework of the IdEX Bordeaux University "Investments for the Future" program / GPR Bordeaux Plant Sciences » to LF, SM. This work was supported by the French National Research Agency (grant no. ANR-19-CE20-0016-02 to DB, LF, SM, FSP, PVD). The authors declare that none of the funding sources was involved in study design, collection, analysis and interpretation of data, writing of the manuscript and decision to submit the work for publication.

## Supplemental information

### SUPPLEMENTAL FIGURES

**Figure S1. Quantification of (h)(VLC)FAMES and sterol moieties released after acidic hydrolysis of AtPM preparation. A,** Representative GC-MS chromatography showing the separation of (h)VLCFAMES moieties from sphingolipid part. Quantification of the different fatty acids (h)(VLC)FAMES detected by GC-MS. h14 and C17 are the internal standard. **B,** Representative GC-MS chromatography showing the separation of unsaturated FAMES with 0-3 double bonds. Quantification of the different fatty acids FAMES detected by GC-MS. C17 is the internal standard. **C,** Representative GC-MS chromatography of sterol moieties. Quantification of the different sterol moieties with cholestanol as internal standards. Cholestanol is the internal standard. Experiments were done 3 times on independent AtPM preparations, mean (± standard deviation) is shown.

**Figure S2. Molecular species analyzed by LC-MS/MS by MRM method of major lipid classes found in AtPM.** AtPM preparations were extracted with the different procedure dedicated to the classes of lipids and injected to LC-MS/MS. Results show the different molecular species of PC, PE, PG, DGDG, MGDG, LCB, LCB-P, CerR, GlcCer, Glysosyl inositol phosphoryl ceramide GIPC series A, Hex(R1)-HexA-IPC and series B Hex-Hex(R1)-HexA-IPC. Analyses were performed 3-4 on independent AtPM preparations. cn:d, with n representing the total number of carbons and d the number of unsaturations, the two fatty acids are represented in parenthesis. A dash (-) between the two fatty acids within the nomenclature means that the regiolocation is not known. Conversely, the use of a slash (/) shows that the regiolocalisation is experimentally established.

**Figure S3. Purification of cauliflower GIPC series A (Hex(R1)-HexA-IPC) to be used as external standards for LC-MS/MS.** GIPC were purified from cauliflower according to (**128**). **A**, Purified GIPC series A (Hex(R1)-HexA-IPC) were analyzed by TLC and **B**, recisely quantification by GC-MS after acidic hydrolysis and derivatization. **C**, Purified GIPC were injected to LC-MS/MS spiked with GlcCer d18:1/c12:0. By estimating that the response of the purified GIPC in LC-MS/MS is similar to that of the GIPC contained in the AtPM samples and using the same internal standard GlcCer d18:1/c12:0, we were able to estimate the amount of GIPC present in the AtPM sample.

**Figure S4.** Unsaturation and chain length of fatty acids for major lipids identified in AtPM.

**Figure S5.** Relationship between the number of proteins identified in the PM proteome of *A*. *thaliana* and the number of available protein entries for *A. thaliana*.

### SUPPLEMENTAL TABLES

**Table S1.** MRM tables used in the lipidomic analysis.

**Table S2.** List of the 3,948 proteins and their corresponding NSAF values obtained from the six fractions analyzed in the current study by using an *E*-value smaller than 10^−5^ as a criterion to assign correct protein identification. Accessions and annotations refer to https://www.araport.org/downloads/Araport11_Release_201606.

**Table S3.** Experimentally know localization of the 3,948 proteins identified in the current study, as inferred from HDA (High Throughput Direct Assay) and IDA (Inferred from Direct Assay) GO experimental evidence codes for cellular components retrieved from SUBA5 and Araport11. Accessions and annotations refer to https://www.araport.org/downloads/Araport11_Release_201606.

**Table S4.** Quantitative distribution of the proteins identified in the three microsomal (Atµ) and plasma membrane (AtPM) fractions.

**Table S5.** *In silico* characterization of the 2,165 proteins corresponding to the core-set of the plasma membrane (PM) proteome of Arabidopsis Col-0 cells when retaining only the proteins co-identified in the three distinct PM replicates analyzed in the current study.

**Table S6.** *In silico* characterization of the 1,759 proteins inferred as experimentally-known PM resident according to HDA (High Throughput Direct Assay) and IDA (Inferred from Direct Assay) GO experimental evidence codes for cellular components searched against SUBA5 and Araport11.

**Table S7** *In silico* characterization of the 406 proteins inferred as experimentally-unknown PM resident according to HDA (High Throughput Direct Assay) and IDA (Inferred from Direct Assay) GO experimental evidence codes for cellular components searched against SUBA5 and Araport11.

**Table S8.** List of the 133 lipid-related proteins identified in the AtPM proteome.

**Table S9.** Comparison of the number of proteins identified over time in the PM proteome of *Arabidopsis thaliana*.

