## Supplementary figures and images for "A combined lipidomic and proteomic profiling of *Arabidopsis thaliana* plasma membrane"

### Figure S1

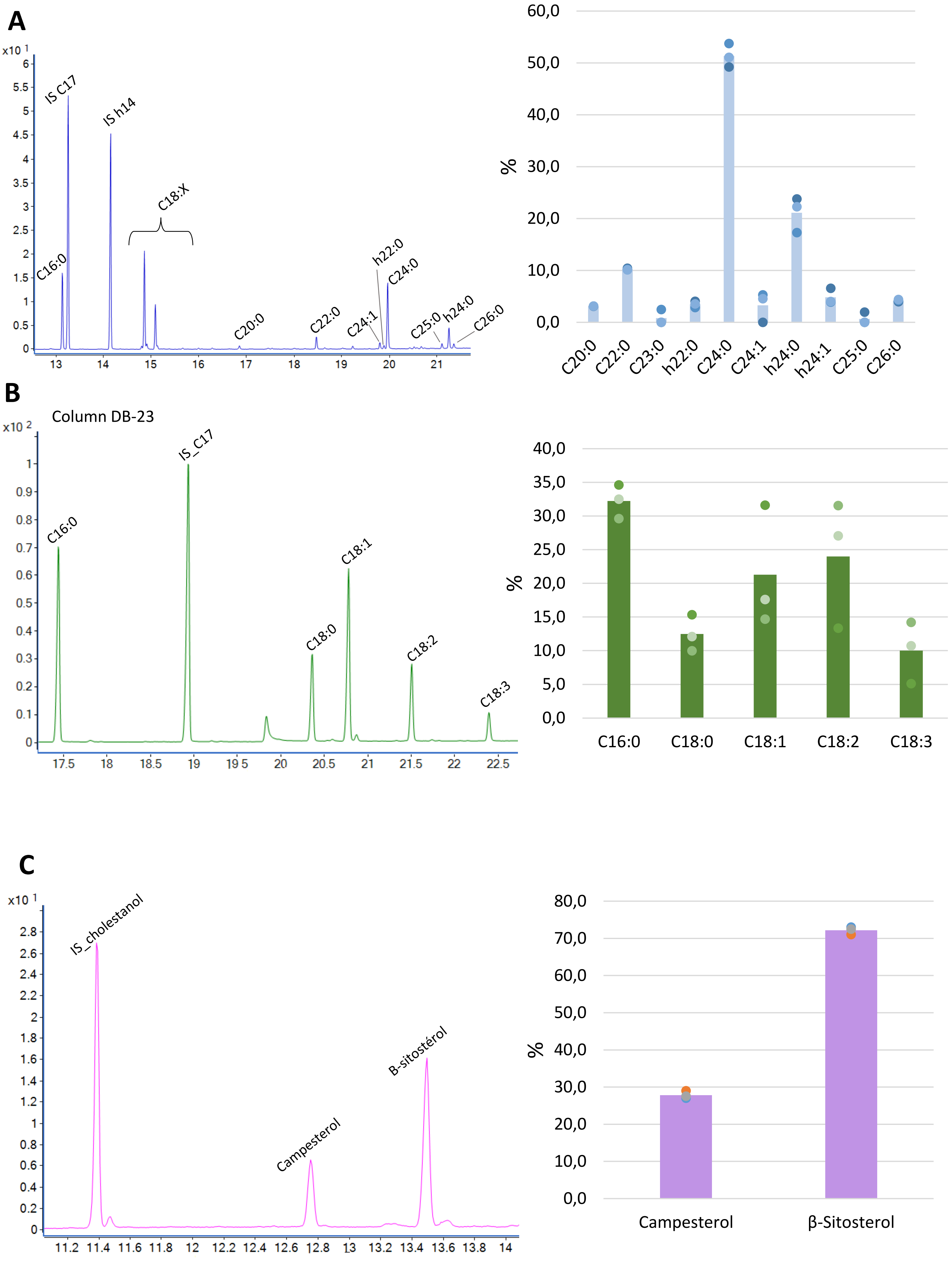

### Figure S2

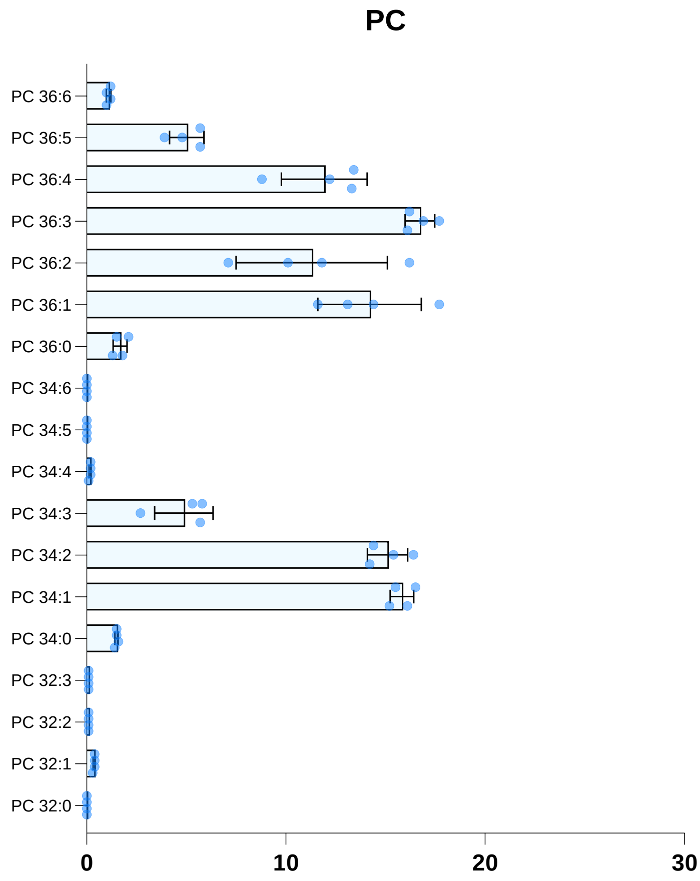

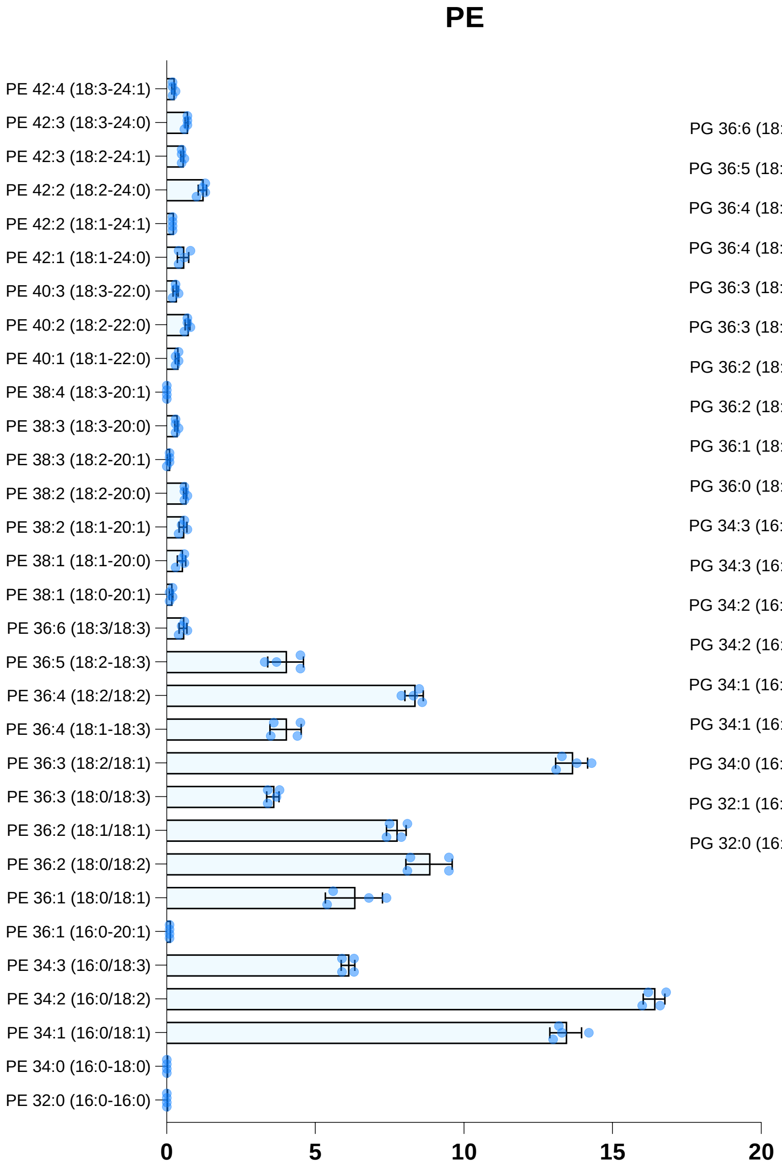

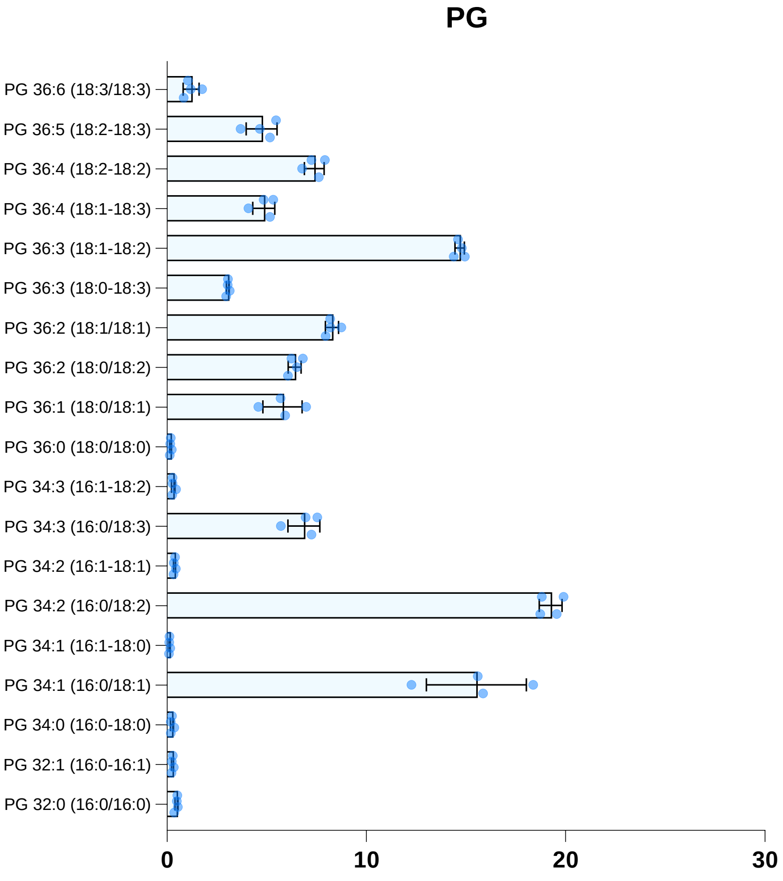

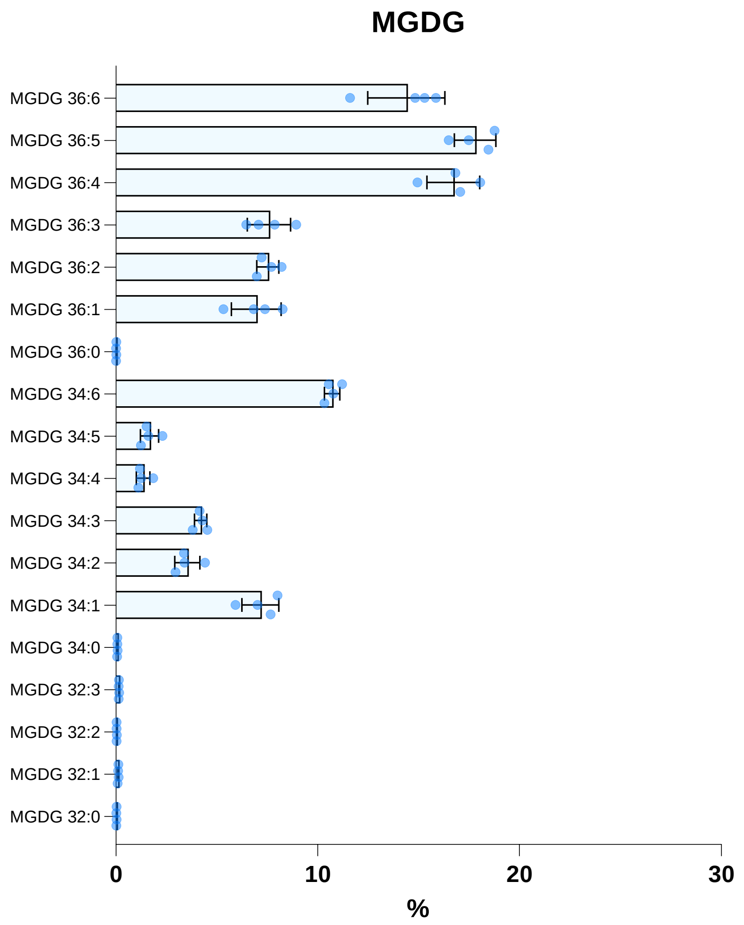

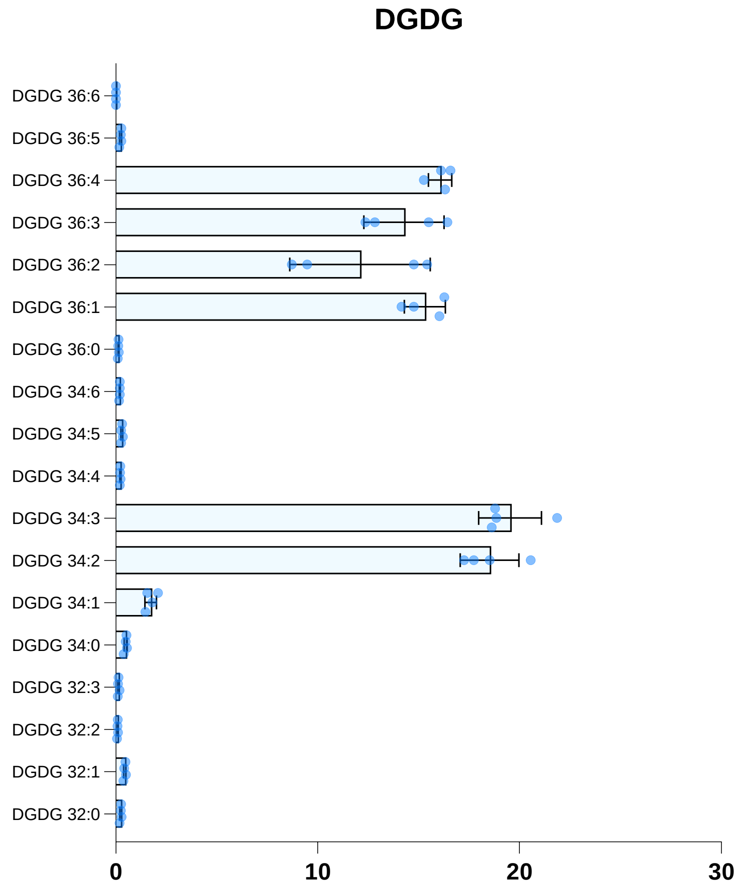

#### Neither LCB, nor LCB-P were detected in AtPM

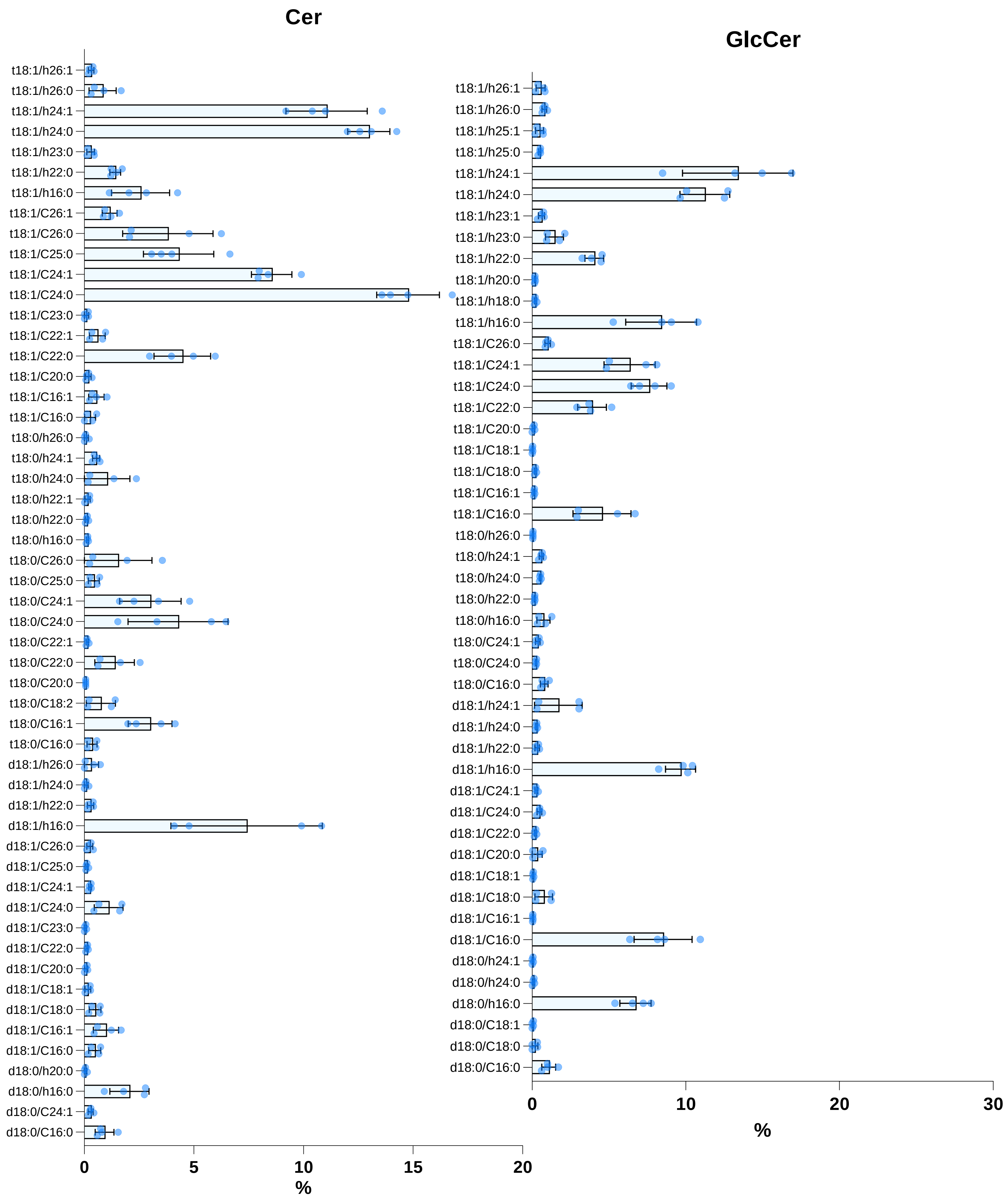

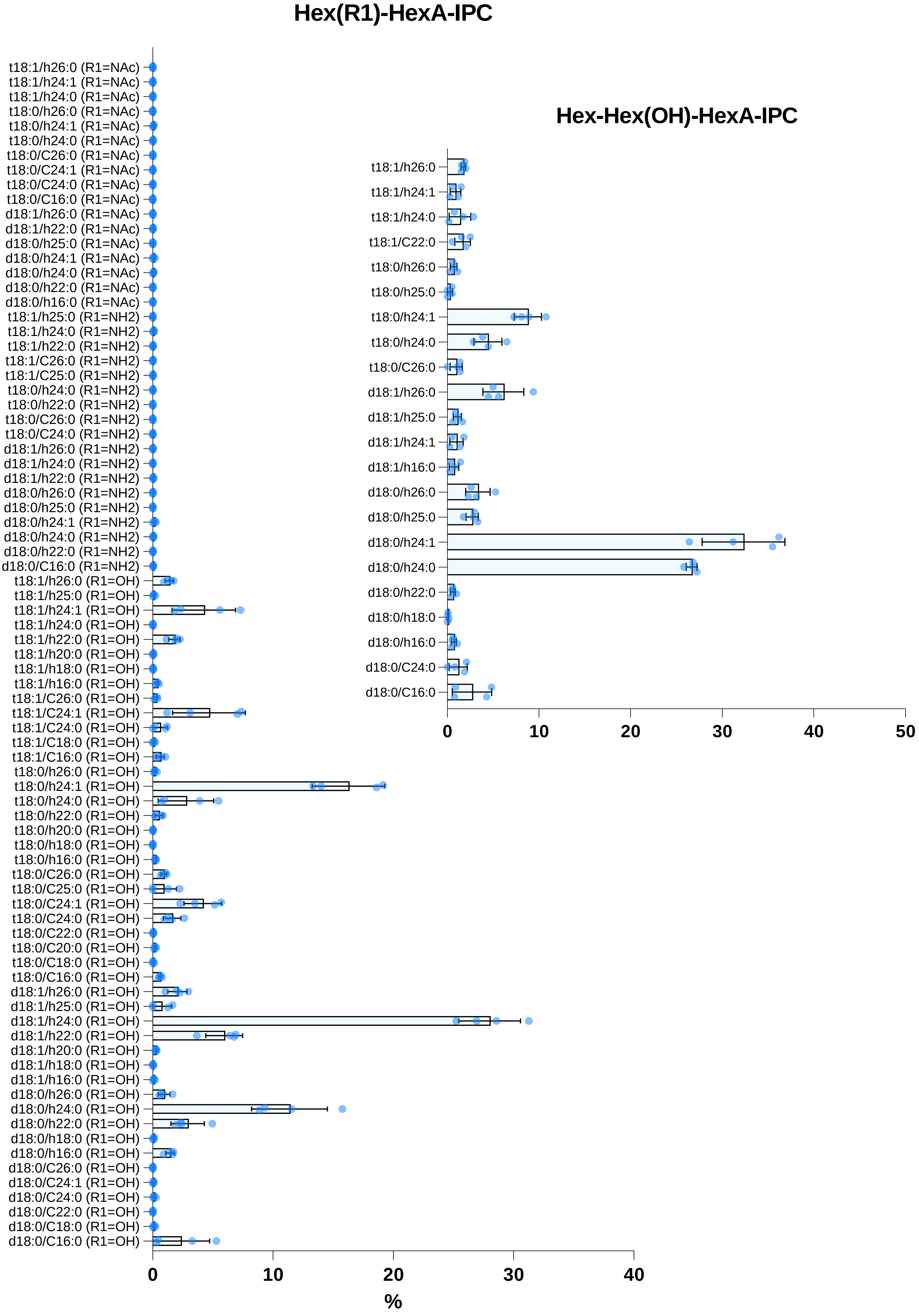

### Figure S3

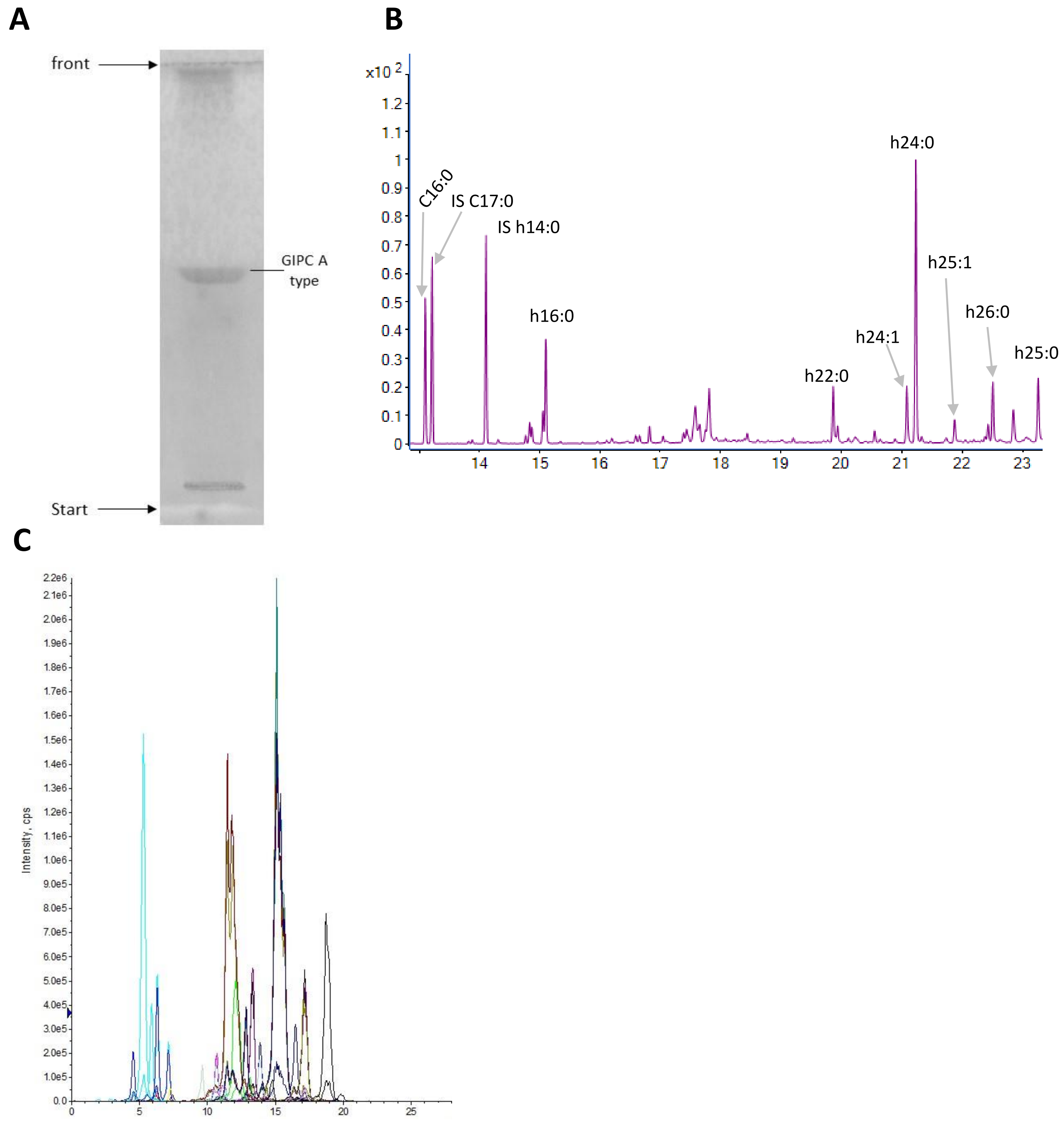

### Figure S4

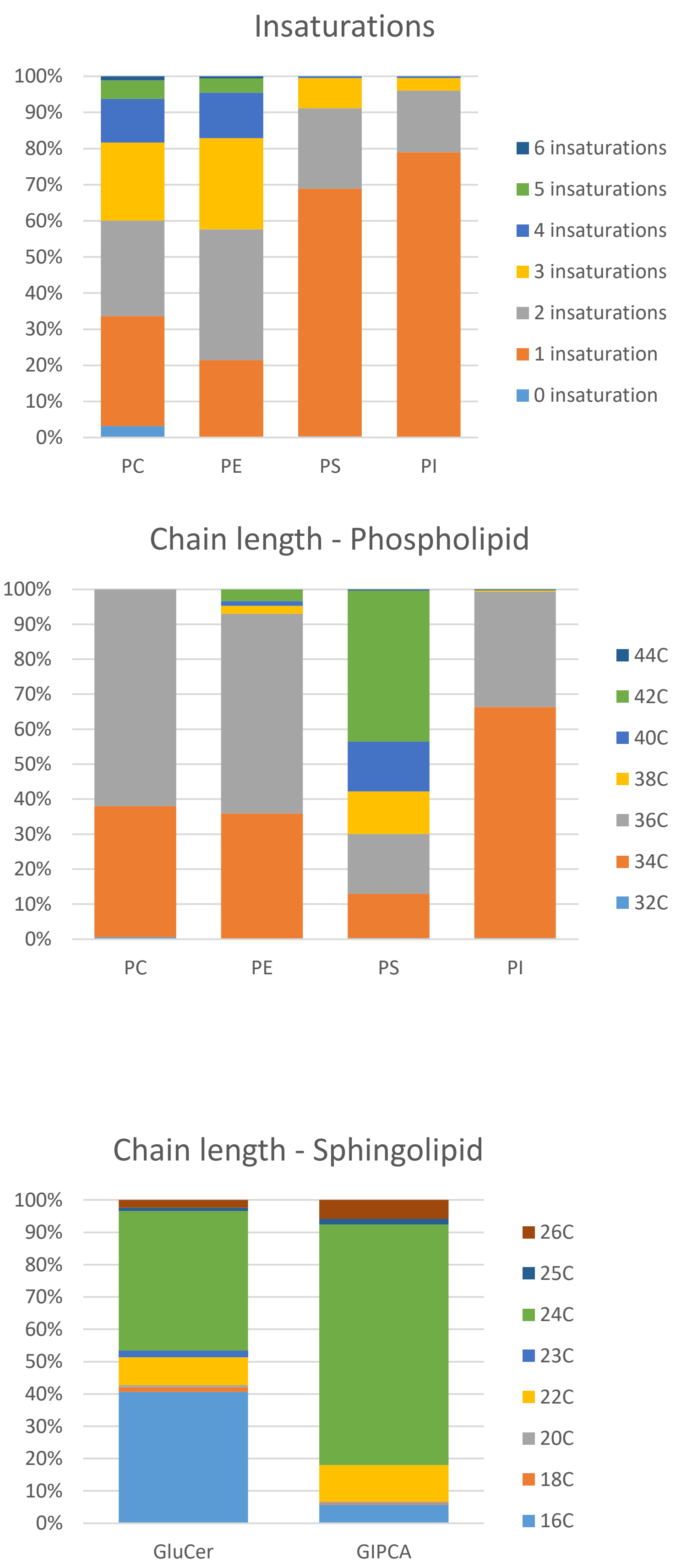

### Figure S5

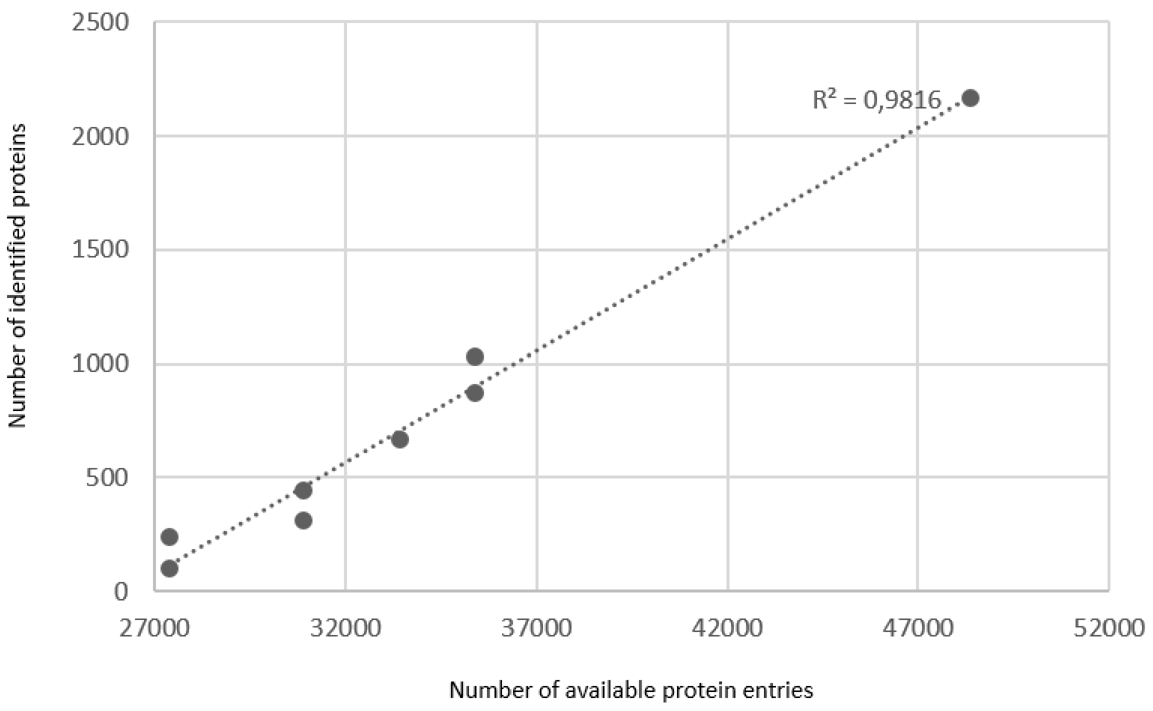
