## Supplementary material for "A combined lipidomic and proteomic profiling of *Arabidopsis thaliana* plasma membrane": Table S4

**Table S4**. Quantitative distribution of the proteins identified in the three microsomal (Atµ) and plasma membrane (AtPM) fractions.

| Number of proteins | Fraction Atµ1 | Fraction Atµ2 | Fraction Atµ3 | Fraction AtPM1 | Fraction AtPM2 | Fraction AtPM3 | Total |
|---|---|---|---|---|---|---|---|
| LC-MS/MS | 3,417 | 3,133 | 3,396 | 2666 | 2,281 | 2,875 | 3,948 |
| | | Atµ common pool: 2,988 | | | AtPM common pool: 2,165 | | |
| IDA/HDA cellular component | 3,293 (96.4%) | 3033 (96.8%) | 3,276 (96.5%) | 2,558 (95.9%) | 2,191 (96.0%) | 2,732 (95.0%) | 3745 (94.8S%) |
| | | Atµ common pool: 2,401 | | | AtPM common pool: 2,040 | | |
| Exclusive IDA/HDA cellular component | 430 (12.6%) | 372 (11.9%) | 437 (12.9%) | 393 (14.7%) | 350 (15.3%) | 441 (15.3%) | 595 (15.1%) |
| | | Atµ common pool: 338 | | | AtPM common pool: 325 | | |
