## Supplementary material for "A combined lipidomic and proteomic profiling of *Arabidopsis thaliana* plasma membrane": Table S9

**Table S9.** Comparison of the number of proteins identified over time in the PM proteome of *Arabidopsis thaliana*.

| Material | Number of proteins identified in the PM proteome | Reference | Number of available protein entries in TAIR |
|---|---|---|---|
| Cell culture | 102 | Marmagne et al. (2004) | 27,416 |
| Leave and petiole | 238 | Alexandersson et al. (2004) | 27,416 |
| Cell culture | 309 | Nelson et al. (2006) | 30,899 |
| Cell culture | 446 | Marmagne et al. (2007) | 30,899 |
| Cell culture | 666 | Zhang and Peck (2011) | 33,410 |
| Seedling | 1,029 | de Michele et al. (2016) | 35,386 |
| Seedling | 873 | Miki et al. (2019) | 35,386 |
| Cell culture | 2,165 | This study | 48,358 |
